# Fin Shape, Asymmetry, and Evolutionary Ecomorphology in Triggerfishes and Filefishes (Superfamily: Balistoidea)

**DOI:** 10.1101/2021.06.29.450391

**Authors:** Andrew B. George, Mark W. Westneat

## Abstract

Triggerfishes and filefishes exhibit a wide range of fin and body morphologies, inhabit many marine habitats, and feed on a variety of benthic and pelagic organisms. Particular morphologies are predicted to provide functional advantages for swimming behaviors that facilitate life in diverse habitats and feeding guilds. Ecomorphological relationships can, in turn, inform evolutionary patterns of morphological convergence. We quantified morphological diversity of 80 balistoid species using geometric morphometrics and assigned each species a primary habitat and feeding mode. Results revealed strong evidence for evolutionary integration among body and fin shapes as well as widespread convergence of both high and low aspect ratio (AR) dorsal and anal fins, the fins that power steady locomotion in these fishes. Dorsal and anal fins were determined to be moderately to highly asymmetrical in most species. Families exhibited considerable overlap in fin and body shapes, but triggerfishes generally exhibited higher AR and more asymmetrical fins than filefishes. Fin asymmetry was not strongly associated with ecology. Planktivorous and offshore-pelagic species exhibited high AR dorsal and anal fins suitable for high endurance swimming performance, while benthic grazing and structured reef species exhibited convergence on low AR median fins more suitable for facilitating maneuverability.

## INTRODUCTION

Fishes inhabit nearly every aquatic ecosystem from the open ocean, structurally complex coral reefs and wave-swept shorelines to freshwater systems including fast-flowing rivers and dark, still cave waters. Understanding which organismal traits allow fishes to successfully inhabit these ecosystems is a general goal of many evolutionary ichthyologists because these patterns can help explain both clade-specific evolutionary trajectories and larger trends between functional traits and ecology. Studies exploring relationships between organismal ecology and morphology, termed ecomorphology (Wainwright and Reilly 1994), have been especially useful for large taxonomic groups because detailed morphological data can be gathered quickly for many species. When conducting an ecomorphological study, it is important that the taxa exhibit convergence in the ecological and morphological traits examined to ensure that any observed evolutionary patterns are not simply a product of shared ancestry. Furthermore, to fully understand ecomorphological relationships, the morphological characters examined must have well-established associations with organismal function or performance, often provided by biomechanical experiments. Consequently, many ecomorphological studies have focused on relationships between cranial morphology and diet (Friedman et al. 2016; Klaczko et al. 2016; McCord and Westneat 2016a; Olsen 2017; Evans et al. 2019; Nicholson and Clements 2021) and locomotor morphology and habitat (Fulton et al. 2001; Yuan et al. 2019). Although most fishes rely on swimming performance for nearly all aspects of their lives (including prey capture), fewer studies (Wainwright et al. 2002) have examined ecomorphological relationships between fish locomotor morphologies and feeding modes.

Triggerfishes (Balistidae) and filefishes (Monacanthidae) in the superfamily Balistoidea are an excellent system for exploring swimming-related ecomorphological relationships due to their wide ranging fin and body shapes, feeding modes, and marine habitats. Balistoid fishes range in body shape from deep-bodied, rounded forms to shallow-bodied, elongate shapes and possess a wide range of fin shapes. Despite all balistoid species using their dorsal and anal fins to power steady swimming with the balistiform swimming mode (Blake, 1978; Wright, 2000), these fishes possess a high degree of morphological diversity in dorsal and anal fin shapes, particularly with regards to fin aspect ratio (AR). Fin AR is a measure of how wing-like a fin is, with high AR fins resembling energetically efficient cruising wings with long leading-edge fin rays, and low AR fins resembling broad, maneuverable wings with uniformly short fin rays (see Figure 1). Dorsal and anal fin shapes range from posteriorly-tapering, high AR forms to rectangular or rounded, low AR forms (Dornburg et al. 2011). Caudal fins, which balistoid fishes use during high-speed steady swimming and burst swimming maneuvers, range from deeply forked to highly convex morphologies (George and Westneat 2019). Laboratory experiments have revealed that balistoid species with high AR dorsal, anal, and caudal fins are associated with increased endurance swimming performance (Wright 2000; George and Westneat 2019).

**Figure 1.**
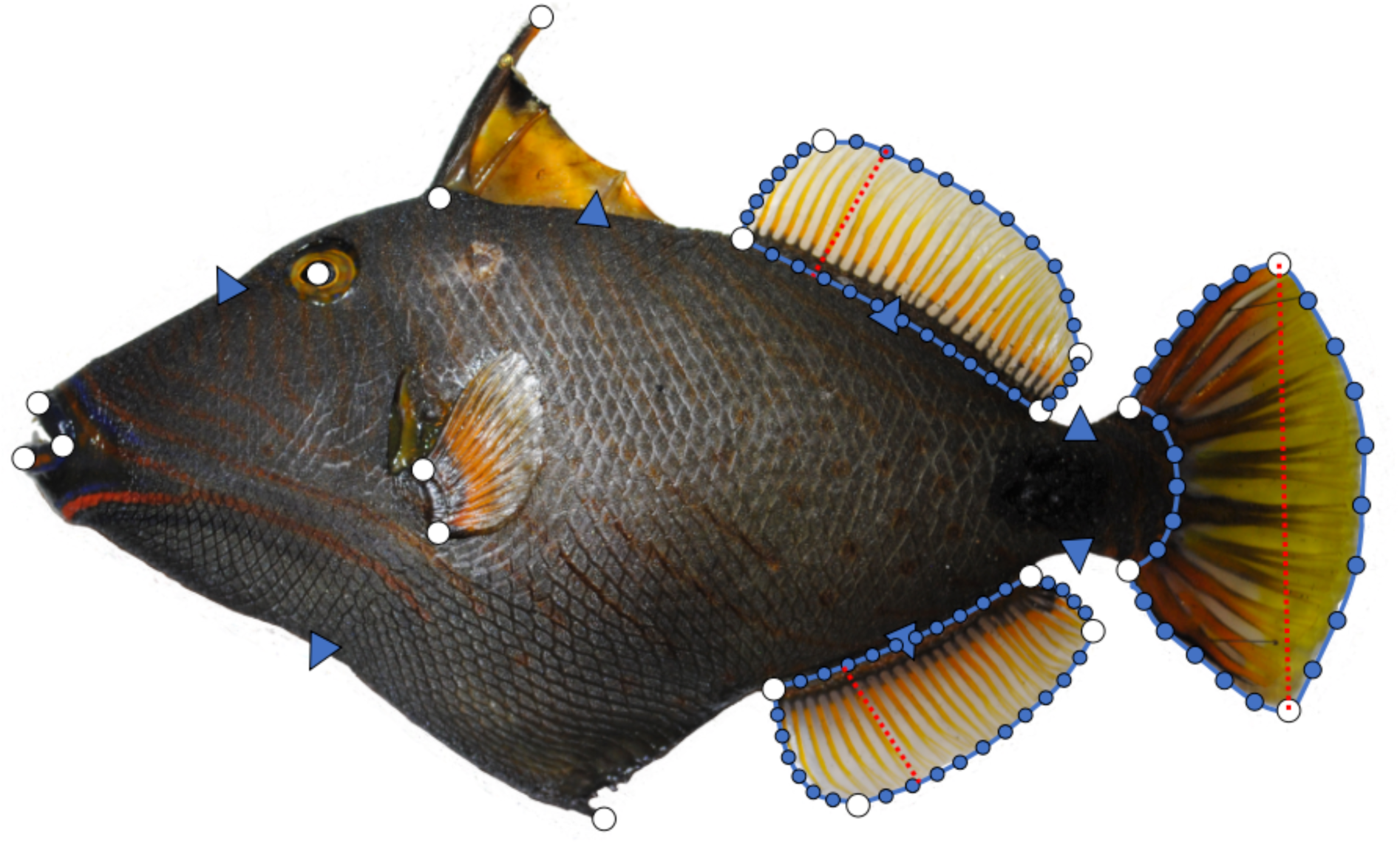
Morphometrics digitization scheme demonstrated on the triggerfish *Balistapus undulatus*. A total of 110 landmarks were placed along the fins and bodies of each fish. White circles represent static landmarks. Blue shapes represent sliding semi-landmarks, with triangles representing landmarks in the body dataset and circles representing subsampled fin landmarks from digitized Bezier curves. Fin area measurements are outlined in blue and span measurements were made along the dotted red lines.

The common ancestor of the superfamily Balistoidea was likely reef associated (Santini et al. 2013), but many extant species throughout the superfamily now occupy a variety of marine habitats including seagrass beds, bare coastal shores, and even the open ocean. Furthermore, many filefish lineages have convergently re-evolved reef associations following ancestral off-reef shifts (Santini et al. 2013). Balistoid fishes also exhibit a variety of feeding modes including benthic grazing on slow moving or sessile organisms, planktivory in the water column, and predation on elusive prey such as cephalopods and other fishes. Previous work has shown that triggerfish cranial morphology is tightly correlated with feeding mode (McCord and Westneat 2016a) and that triggerfish dorsal and anal fins exhibit high degrees of morphological integration across species (Dornburg et al. 2011). However, little remains known about relationships between balistoid diet and habitat ecologies and swimming-related morphologies.

In the present study, balistoid fin and body geometric morphometrics are used to examine patterns of morphological evolution of functionally informative traits and the relationships between these traits and balistoid ecology. Based on observation of extant species, we predicted a high degree of convergence in dorsal, anal, and caudal fin shapes across the superfamily. Due to the strong functional relationships documented between balistoid fin morphology and swimming performance (George and Westneat 2019), we hypothesized that fin shape convergence was associated with feeding mode and habitat use ecology. Specifically, we tested the hypotheses that planktivorous and pelagic balistoid fishes possess higher aspect ratio fins than fishes in other feeding mode and habitat categories. This prediction is based on the high percentage of time fishes in these groups spend swimming in the water column, thus benefitting from the performance (George and Westneat 2019) and hydrodynamic (Wright 2000) advantages provided by high AR fins. Conversely, low AR fins were hypothesized to be associated benthic grazers and species that live in structured reef environments because low AR fins are often associated with high performance burst swimming and maneuverability (Walker and Westneat 2002), characteristics likely to facilitate predator avoidance in structurally complex habitats and grazing performance, respectively. Similarly, deep ventral keels and long dorsal spines were expected to be associated with fishes living in structured reef environments because these mobile elements can be used to avoid predation by safely locking fishes into reef crevices. Finally, recent work (George and Westneat, 2019) and initial observations of balistoid body plans suggest that dorsal and anal fin shapes and positions are not symmetrical across the dorsal-ventral body axis in many species, and we set out to test the hypothesis that the degree of fin asymmetry may also be associated with dietary traits and habitat ecology.

## MATERIALS AND METHODS

### Species Selection, Habitat Use, and Feeding Traits

We examined the 80 balistoid species in the most recent molecular phylogeny of the superfamily (McCord and Westneat 2016b), including 32 triggerfish (Balistidae) species and 48 filefish (Monacanthidae) species, representing 54% of the 150 described balistoid species (76% of triggerfishes and 45% of filefishes). This time-calibrated phylogeny was used for all phylogenetically-informed statistical analyses in this study.

We assembled a trait matrix of primary habitat use and feeding mode for all 80 species based on published literature. A literature survey of 99 articles, books, and encyclopedias (see Table S1 and supplemental methods for details) was performed to assign balistoid species into 7 habitat categories (coastal bare bottom, structured reef, open reef, open ocean pelagic, open ocean demersal, open ocean rafting, and seagrass/ weedy) and 4 feeding mode categories (benthic grazers, elusive prey feeders, planktivores, and a benthic grazing + planktivory category). When habitat and feeding data were not available from primary sources, the online database *FishBase* (Froese and Pauly 2019) was used to classify remaining species into habitat (n = 4) and feeding groups (n = 1).

### Fin and Body Shape Quantification

Morphological variation of fin and body shapes among the 80 balistoid species was quantified using photos of 570 fish specimens acquired through online museum databases, publications, online fish blogs, and physical museum specimens (see Table S2 for photograph sources). Each species was represented by an average of 6.6 specimens (range = 1-20). As morphometric quality of photos was prioritized over quantity, 4 rare species (*Eubalichthys mosaicus*, *Arotrolepis filicauda*, *Paramonacanthus oblongus*, and *Thamnaconus arenaceus*) were represented by 1 specimen, and 7 species were represented by 2 specimens. All other species were represented by 3 or more specimens.

A total of 110 landmarks were digitally placed on each photo using the R package *StereoMorph* (Olsen and Westneat 2015) for geometric morphometric analyses, following the methods of George & Westneat (2019) with the exception of one additional landmark on the distal tip of the first dorsal spine in the present study (Figure 1). If any structures of a specimen were visibly damaged, those structures were not digitized or included in subsequent analyses. Some dorsal and anal fins (n=96/336 and 83/329, respectively) in otherwise good condition had preservation artifacts leading to inconsistent fin positions between specimens. These preservation artifacts were corrected using the Mac App *FinRotate* (https://github.com/mwestneat/FinRotate) as described in George & Westneat (2019) (Table S3). All dorsal spine tip landmarks were rotated with *FinRotate* so that a consistent 45 degree angle was formed between the dorsal spines and the anterior-posterior body axis. Similarly, the positions of un-extended ventral keel landmarks were estimated using the *estimate.missing* function in the *geomorph* R package (Adams et al., 2017) following the methods of George & Westneat (2019).

Landmarks were subdivided into five separate morphometric datasets for independent geometric morphometric analyses: full shape (all landmarks), body only, dorsal fin only, anal fin only, and caudal fin only following the methods of George & Westneat (2019). The additional dorsal spine landmark in the present study was treated as a static landmark and included in the full shape dataset only. Following Procrustes-alignment, species-average shapes for each morphological dataset were calculated, and allometric relationships between log-transformed species centroid size and shape were assessed using the *procD.pgls* function in *geomorph* based on 1,000 permutations. As no significant allometric relationships were detected in the dorsal and caudal fin datasets (F = 2.3366, P = 0.0847; F= 1.2922, P = 0.244, respectively), no adjustments to these morphometric data were required, and principal components analysis (PCA) was conducted on each of these species-averaged, Procrustes-aligned geometric morphometric dataset. Resulting principal component (PC) scores along the two most significant axes of inter-specific variance (PCs 1 and 2) were extracted for each species for use in backtransform morphospace visualization (see Olsen, 2017) and subsequent statistical analyses. Allometric relationships were detected in the full shape, body, and anal fin datasets (F = 2.6612, P = 0.0119; F = 3.2564, P = 0.0175; F = 2.8017, P = 0.0355, respectively). To remove covariation between size and shape, residuals from *procD.pgls* analyses were used to extract allometry-free full shape, body, and anal fin shapes for each species. These size-adjusted shapes were used in all subsequent analyses for these morphological datasets. Final sample sizes (species, specimens) for each morphological unit are: body (80, 530), dorsal fin (76, 336), anal fin (77, 329), caudal fin (76, 254), and full shape (74, 175).

Finally, dorsal, anal, and caudal fin aspect ratios (AR) were calculated for each specimen using the rotated fin landmarks and the equation:

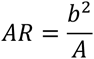

where *b* is the maximum span of each fin (red dotted lines in Figure 1) and *A* is the surface area of the fin (blue lines in Figure 1) (George and Westneat 2019). Species-average dorsal, anal, and caudal fin ARs were then calculated.

### Evolutionary Morphological Integration

Two methods of pair-wise morphological integration tests were conducted between each geometric morphometric dataset (body, dorsal fin, anal fin, and caudal fin) in a similar approach to that used by Dornburg et al. (2011). First, pair-wise phylogenetic integration tests were conducted between geometric morphometric datasets using the *phylo.integration* function in *geomorph*. In a pair-wise context, the *phylo.integration* test uses two-block partial least squares (PLS) analysis to quantify the degree of morphological covariation (integration) among species in a phylogenetic context. Pairwise linear regressions were used to assess significance of correlation between primary phylogenetic partial least squared (pPLS) axes of each morphological unit. Second, separate pair-wise phylogenetic generalized least squares (PGLS) regressions were conducted between all combinations of the first two principal component (PC) axes of each geometric morphometric dataset using the *pgls* function in the *caper* R package (Orme et al. 2013). See the Supplemental Methods for a detailed description of the differences between these two integration tests.

### Phylogenetic ANOVA and Convergence Testing

Two methods were used to assess relationships between balistoid morphology and ecology. First phylogenetic ANOVA tests were conducted between fin ARs and significant PC axes (PCs 1 and 2) of each geometric morphologic dataset (full shape, body, dorsal fin, anal fin, and caudal fin) and the habitat and feeding mode categories using the *phylANOVA* function in the R package *phytools* (Revell 2012). Statistical significance of phylogenetic ANOVA tests were based on 10,000 simulations. Pair-wise t-tests were then conducted to identify ecotypes (habitat or feeding groups) exhibiting significantly different morphologies from one another (see Supplemental Methods for further details). Second, the *convevol* R package (Stayton 2018) was used to test for morphological convergence among species in the same feeding mode and habitat use groups. Specifically, C1 convergence (the proportion of the maximum ancestral distance between focal species that has been closed over evolutionary time) of each morphological dataset was assessed for species in the open-ocean pelagic, structured reef, planktivorous, and benthic grazing ecotypes to test ecomorphological hypotheses. Statistical significance of convergence tests were based on 500 simulations. All p-values resulting from phylogenetic ANOVA post-hoc pair-wise t-tests and from morphological convergence tests were adjusted for multiple tests using the “holm” method (Holm 1979). Statistical significance was determined as p <0.05.

### Dorsal and Anal Fin Symmetry

Dorsal and anal fin symmetry was assessed in all 287 specimens with both dorsal and anal fin data, for 76 species. Median fin shape symmetry was quantified using three morphological ratios: 1) Dorsal fin AR: Anal fin AR, 2) Dorsal fin area: Anal fin area, and 3) Dorsal fin base length: Anal fin base length. Symmetry analyses requiring perfectly positioned fins in the same photograph were limited to the 175 specimens (74 species) included in the full shape dataset. Median fin position and angle symmetry was assessed by comparing the position of the first dorsal and anal fin rays along the anterior-posterior axis and by comparing the angles of dorsal and anal fin attachment, respectively. We used the *bilat.symmetry* function in *geomorph* to comprehensively measure symmetry/ asymmetry using ANOVA tests between dorsal and anal fin landmarks. Two separate *bilat.symmetry* tests were conducted to assess the degree of directional symmetry (Klingenberg et al. 2002). First the null hypothesis of matching symmetry was tested by flipping the anal fin coordinates over the body axis (to remove the mirror-image effect) and Procrustes aligning the dorsal and anal fin landmarks together. This process removes information about translation, rotation, and scaling between dorsal and anal fins, and thus only tests the degree of symmetry between the shapes themselves, regardless of position on the body. Next, the null hypothesis of object symmetry (symmetry across a “midline” axis) was tested. Object symmetry takes the position, size and orientation of the dorsal and anal fins into account while assessing the level of symmetry between the two fins (Klingenberg et al. 2002). The final asymmetry test involved calculating the Procrustes distance (the square root of the sum of squared differences in the positions of the landmarks of two shapes) between Procrustes-aligned dorsal fins and flipped anal fins in order to measure the total spatial transformation shape differences between the fins without taking size or orientation along the body axis into account.

### Fin Ancestral State Estimations

Each species was binned into one of four categorical aspect ratio groups (low, medium, high and very high) using a modified gap coding method (Archie 1985) to allow for relatively even group sizes while ensuring large gaps between species at the inter-group boundaries. Individual AR ancestral state estimations were then conducted for each fin using the *ace* function in the R package *ape* (Paradis et al. 2004) using an equal rates, maximum likelihood model. Evolution of fin asymmetry was explored using ancestral state estimation of the Procrustes distances between dorsal and anal fin shapes. Log-transformed Procrustes distance was treated as a continuous trait, and ancestral states were estimated using the *contMap* function in *phytools*. All statistical analyses were carried out in R version 4.0.2 (R Core Team 2020).

## RESULTS

### Evolution of Fin Aspect Ratios

The balistoid fishes included in this study exhibit a wide range of fin shapes. Dorsal fin aspect ratio (AR) ranged by an order of magnitude from 0.228 in the filefish *Brachaluteres jacksonianus* to 2.49 in the triggerfish *Canthidermis sufflamen*. Similarly, anal fin AR ranged from 0.250 in the filefish *Aluterus scriptus* to 2.45 in *C. sufflamen.* Balistoid fishes also exhibited a wide range of caudal fin AR from 0.742 in the filefish *Aluterus heudelotii* to 4.29 in the triggerfish *Abalistes stellatus*. Triggerfishes tend to have higher dorsal fin AR (range = 0.365 – 2.486, mean = 0.852) and anal fin AR (range = 0.379 – 2.453, mean = 0.786) than filefishes (dorsal fin range = 0.228 – 1.33, mean = 0.477; anal fin range = 0.250 – 1.16, mean = 0.471) (t = 3.72, df = 39.7, p = 0.0006176; t = 3.74, df = 38.8, p = 0.0005917 for dorsal and anal fins, respectively). Balistoid fishes exhibit even larger differences in caudal fin AR between families (t = 9.75, df = 63.63, p = 3.05e^-14^) with triggerfishes generally exhibiting much higher caudal fin AR (range = 2.07 – 4.29, mean = 2.99) than filefishes (range = 0.74 – 2.75, mean = 2.01).

Ancestral state estimations of dorsal, anal, and caudal fin AR revealed multiple convergence events on high, medium, and low AR in all three fin groups and interesting differences in the evolution of these traits between families (Figures 2 and 3). Despite these convergence events, closely related taxa tend to exhibit similar fin ARs, and all three fin ARs showed high levels of phylogenetic signal (K = 0.36, P = 0.001; K = 0.34, P = 0.001; and K = 0.25, P = 0.001, for dorsal, anal, and caudal fins, respectively). Dorsal and anal fin AR are highly correlated (PGLS: P = 2.20e^-16^, multiple R^2^ = 0.775), but caudal fin AR is not correlated with dorsal or anal fin AR (PGLS: P = 0.0787, multiple R^2^ = 0.0418; P = 0.148, multiple R^2^ = 0.0284, respectively).

**Figure 2.**
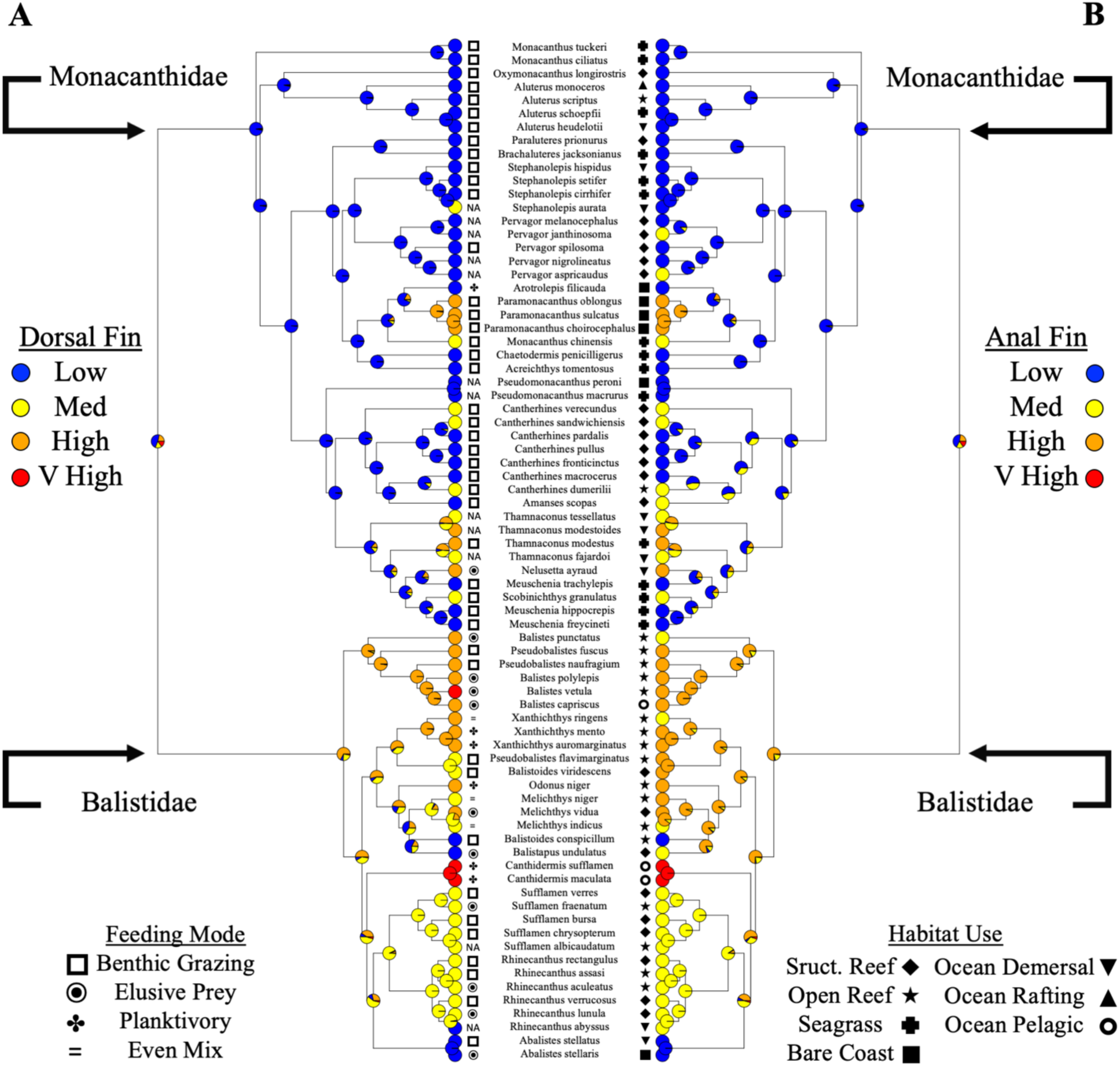
Dorsal and anal fin aspect ratio ancestral state estimations with ecotype labels. (A) Dorsal fin. Low = 0.23 – 0.45, Med = 0.47 – 0.67, High = 0.76 – 1.33, Very High = 2.08 – 2.49. (B) Anal fin. Low = 0.25 – 0.44, Med = 0.46 – 0.66, High = 0.68 – 1.20, Very High = 2.07 – 2.45. Colored pie charts represent the likelihood of each fin aspect ratio state at each ancestral node based. Tip colors depict the measured states of extant species. Black and white tip symbols indicate the feeding mode and habitat use ecotype of each species. “NA” indicates a lack of ecotype data.

**Figure 3.**
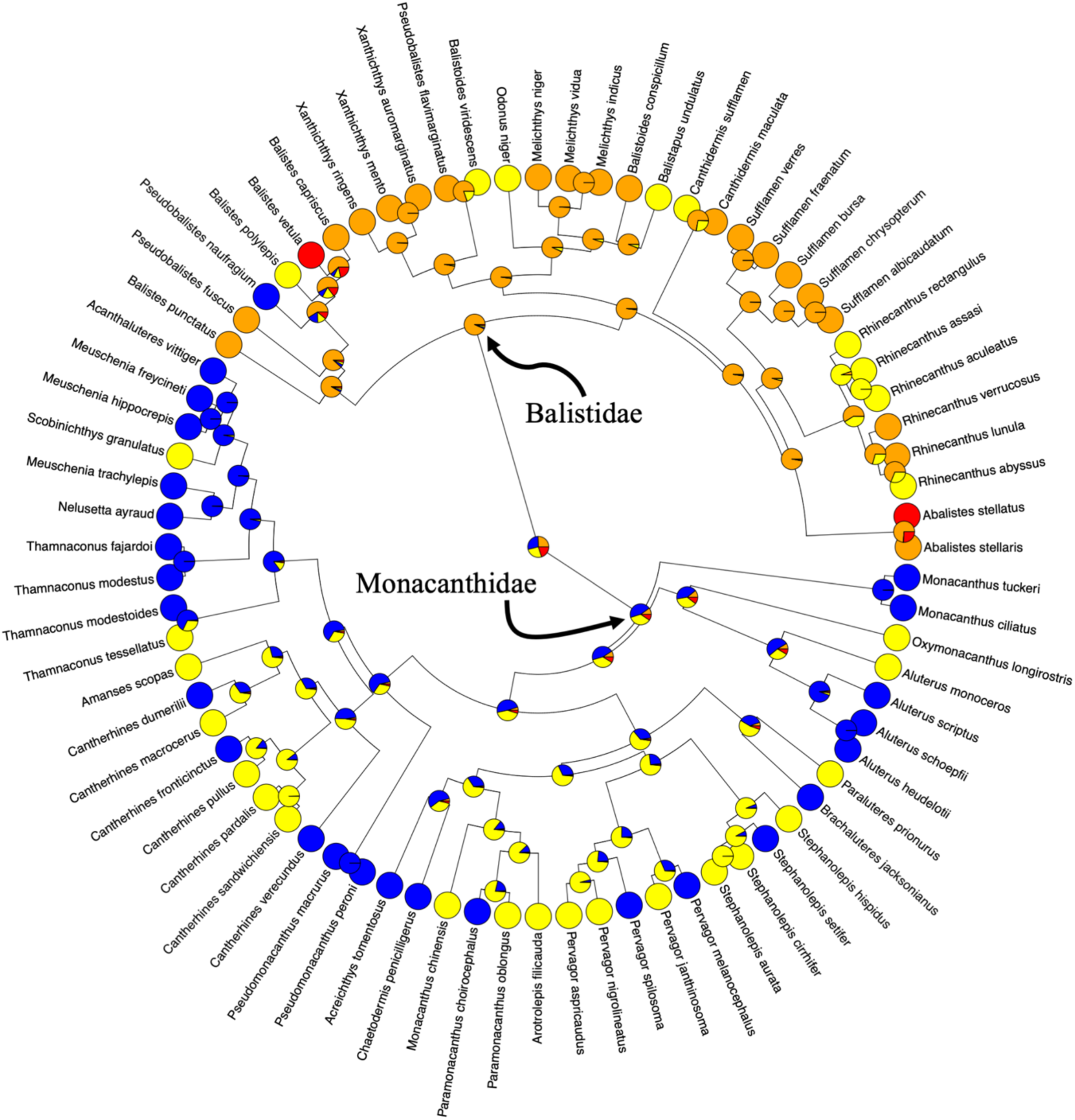
Caudal fin aspect ratio ancestral state estimation. Colored pie charts represent the likelihood of each caudal fin aspect ratio state at each ancestral node. Tip colors depict the measured states of extant species. Low (blue) = 0.74 – 2.09, Med (yellow) = 2.14 – 2.75, High (orange) = 2.81 – 3.61, Very High (red) = 4.14 – 4.29.

The triggerfish common ancestor likely possessed high AR dorsal, anal, and caudal fins, and most extant triggerfishes possess high or medium AR fins (Figures 2 and 3). The highest AR dorsal, anal, and caudal fins were found within the Balistidae. Very high AR dorsal fins likely evolved once each in *Balistes vetula* and the common ancestor to the *Canthidermis* genus, and very high AR caudal fins likely evolved once in *B. vetula* and once in *Abalistes stellatus*. Very high AR anal fins are found only in the *Canthidermis* genus. Medium AR anal fins exhibit the highest degree of convergent evolution within the Balistidae, likely evolving 5 times independently (Figure 2B). These convergence events include the common ancestor of the *Sufflamen + Rhinecanthus* clade, and four individual species, *Balistapus undulatus, Melichthys vidua, Xanthichthys ringens,* and *Balistes punctatus*.

A few triggerfish species possess low AR dorsal, anal, and caudal fins. Low AR dorsal and anal fins likely convergently evolved a few times each within Balistidae (Figure 2). Specifically, low AR dorsal fins likely evolved in the common ancestor of the species pair *Balistoides conspicillum* + *Balistapus undulatus*, in the common ancestor of the *Abalistes* genus, and in *Rhinecanthus abyssus*. Low AR anal fins likely evolved twice within Balistidae, once in the *Abalistes* common ancestor, and once in *B. conspicillum*. Only one triggerfish species, *Pseudobalistes naufragium*, possesses a low AR caudal fin.

The common ancestor of the Monacanthidae, on the other hand, shows a maximum likelihood estimation of low AR dorsal and anal fins and a low-to-medium AR caudal fin. Most extant filefishes possess low AR dorsal and anal fins. As in the Balistidae, medium AR dorsal and anal fins evolved independently many times in the filefishes. Interestingly, 8 filefish species exhibit medium AR dorsal fins and 10 filefish species possess medium AR anal fins, and in both cases none are sister species. Consequently, it appears that medium AR dorsal and anal fins have evolved independently up to 8 times each. High AR dorsal and anal fins most likely evolved 4 times each in monacanthids, once in the common ancestor to the *Paramonacanthus* genus, twice within the *Thamnaconus* genus, and once in *Nelusetta ayraud*. Caudal fin evolutionary trends are more complex in filefishes due to an ambiguous ancestral state, with similar likelihoods of low and medium AR. If the ancestral state was a medium AR caudal fin, then low AR fins convergently evolved up to 9 times, while if the ancestral state was low AR, then medium AR fins evolved up to 7 times independently. However, none of the extant filefishes examined in this study exhibited high or very high AR caudal fins, making it highly likely that the filefishes ancestor possessed a low or medium AR caudal fin.

Results show that the common ancestor of balistoid fishes likely possessed low AR dorsal and anal fins, and that low, medium, and high AR dorsal and anal fins have convergently evolved multiple times throughout the superfamily (Figure 2). The ancestral state of the balistoid caudal fin was not clearly resolved, however triggerfishes and filefishes likely diverged in caudal fin shape shortly following the split between the families, with the triggerfish common ancestor very likely possessing a high AR caudal fin and the filefish common ancestor likely possessing a low or medium AR caudal fin.

### Dorsal and Anal Fin Symmetry

Balistoid dorsal and anal fins exhibited asymmetry along multiple shape and position axes. Although dorsal and anal fin ARs are highly correlated across species, a linear regression between dorsal and anal fin ARs for all 287 individuals (representing 76 species) in which both dorsal and anal fin AR data were available revealed that dorsal and anal fin ARs are not symmetrical within individuals (slope = 0.80). Twelve species possess dorsal and anal fin ARs binned into different AR groups (Figure 2) and only 40% of individuals exhibit the same dorsal and anal fin AR when rounded to one decimal place, demonstrating the widespread asymmetry of this trait. Furthermore, although exactly half the species exhibit higher AR dorsal fins than anal fins, while the other half show the reverse pattern, all species possess longer dorsal than anal fins with the exception of the filefish genus *Aluterus*, which all possess longer anal fins than dorsal fins. A two-sample t-test indicated that balistoid dorsal fin bases, on average, are significantly longer than anal fin bases (t = 2.3098, df = 344.35, P = 0.02149). Additionally, over 90% of individuals exhibited anal fins that begin posteriorly to their dorsal fins. A linear regression also revealed deviations from symmetry in terms of fin angles of attachment (slope = 0.54). Conversely, dorsal and anal fin areas are relatively symmetrical (slope = 0.9397). The discovery of asymmetrical fin aspect ratios, base lengths, and angles of attachment despite highly symmetrical fin areas, suggests that the dorsal and anal fin *shapes*, *positions*, and *orientations*, but *not sizes* have evolved asymmetrically.

Furthermore, directional symmetry tests between dorsal and anal fin landmarks revealed significant matching asymmetry (F = 34.505, P = 0.001) and object asymmetry (F = 318.89, P = 0.001), providing strong evidence for asymmetry in both the shape (primarily fin length) and orientation (primarily angle of attachment) of balistoid dorsal and anal fins. Procrustes distance between aligned dorsal fins and flipped anal fins revealed that triggerfishes generally possess more asymmetrical fins than filefishes (t = 5.4256, df = 167.44, P = 1.995e^-7^). Ancestral state estimations of these Procrustes distances revealed convergent evolution towards asymmetrical median fins throughout the Balistoidea, with the triggerfish ancestor possessing moderately asymmetrical fins, and filefishes evolving from an ancestor with more symmetrical fins (Figure 4).

**Figure 4.**
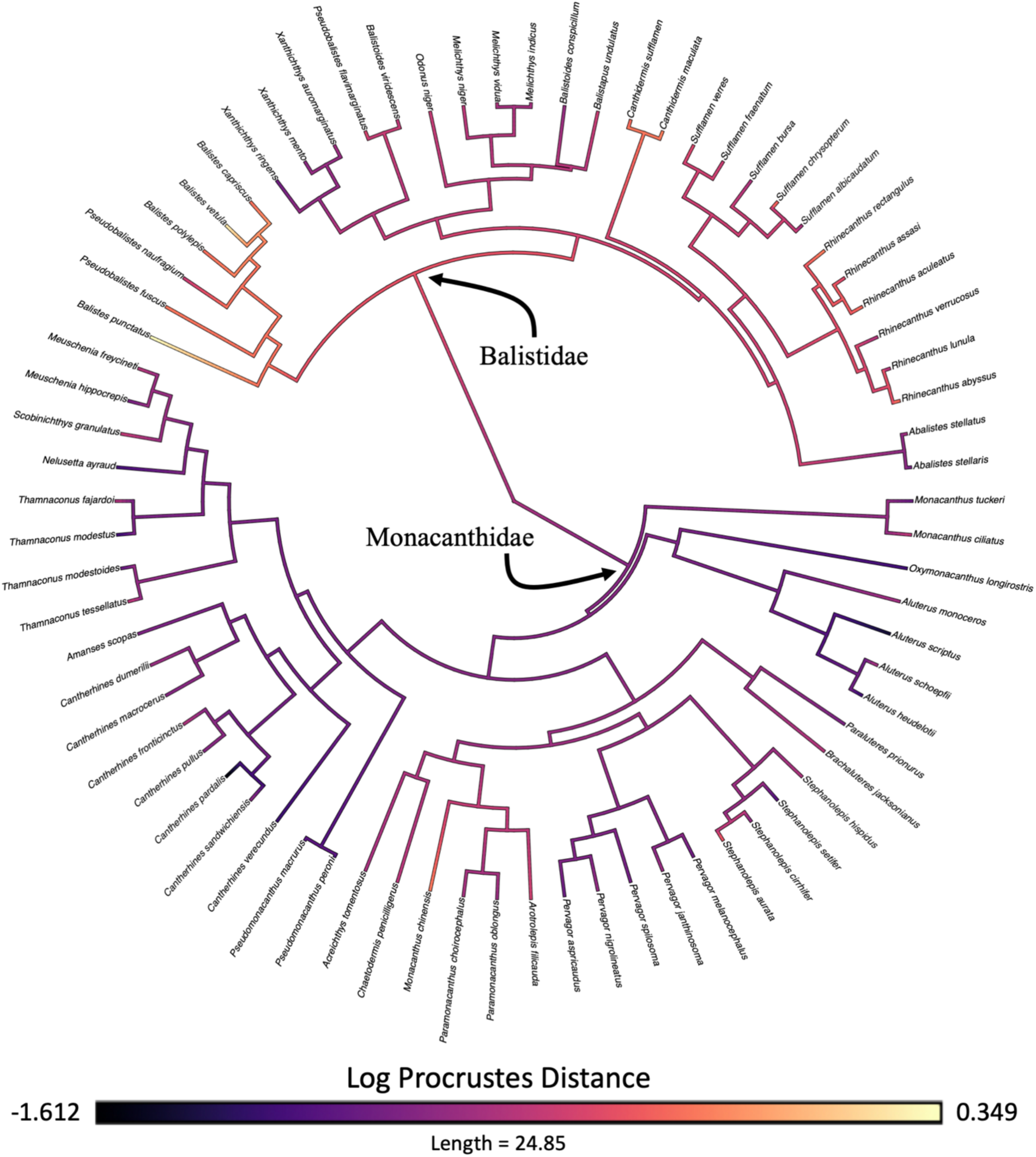
Ancestral state estimation of dorsal and anal fin asymmetry measured as log Procrustes distance. Light and dark colors represent high and low levels of fin asymmetry, respectively. Tip colors represent measured fin asymmetry and nodes colors represent the most likely asymmetry states as determined by the ancestral state estimation model.

### Geometric Morphometrics

The primary axis of variation in the full shape dataset (PC1: 37%) describes changes in the overall length-depth ratio of the fishes with long, thin fishes occupying areas of low PC1 morphospace and short, deep-bodied fishes occupying areas of high PC1 morphospace (Figure 5A). The second axis of variation (PC2: 18%) encompasses differences in dorsal and anal fin ARs, caudal fin shape, and dorsal spine length. Fishes in areas of high PC2 morphospace have long dorsal spines, convex caudal fins, and low AR dorsal and anal fins (Spearman’s rank correlation rho = -0.5191, P = 3.07e^-6^; rho = -0.3957, P = 0.0005336, respectively). Conversely, fishes occupying areas of low PC2 morphospace have high AR, posteriorly tapering dorsal and anal fins, and forked or concave caudal fins. Families exhibit considerable overlap in full shape morphospace, however monacanthids tend to cluster towards high PC2 scores (low AR fins) with the exceptions of *Paramonacanthus oblongus*, *P. choirocephalus*, and *Nelusetta ayraud,* which exhibit higher AR fins and group with many triggerfishes. Filefishes span the full range of PC1 scores, while triggerfishes cluster in central PC1 space defined by intermediate body depths.

**Figure 5.**
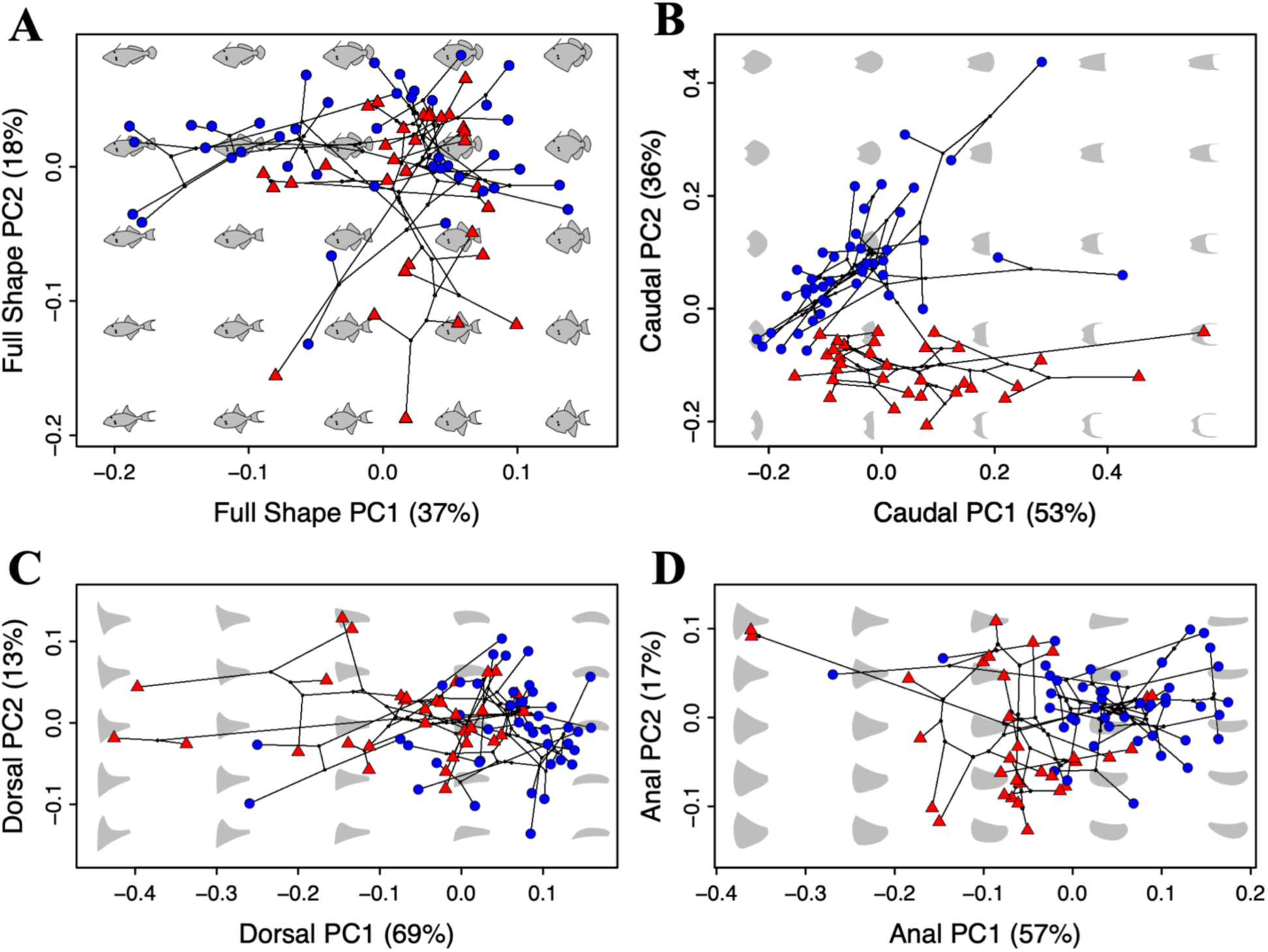
Balistoid fish fin and body shape diversity. (A) Full shape. (B) Caudal fin shape. (C) Dorsal Fin Shape. (D) Anal Fin Shape. Colored shapes represent each species’ position in morphospace with triggerfishes (Balistidae) represented by red triangles and filefishes (Monacanthidae) represented by blue circles blue. Gray shapes represent theoretical backtransformed shapes corresponding to morphologies representative of each location in morphospace. Black lines depict the balistoid phylogeny transformed into each morphospace to facilitate inspection of evolutionary shape change trajectories. Bifurcation points along these lines represent theoretical positions (and morphologies) of each ancestral node.

Species differ in caudal fin morphospace primarily by the shape of the distal edges of their fins (PC1: 53%) with convex caudal fins occupying areas of low PC1 morphospace and forked or concave caudal fins occupying areas of high PC1 morphospace (Figure 5B). Both families tend towards low PC1 scores (convex fins) with just a few members of each family in high PC1 morphospace (concave fins). Caudal fin PC2 (36%) describes the length-depth ratio of the fins and is associated with caudal fin AR (Spearman’s rank correlation rho = -0.9578, P = 2.20e^-16^). Balistoid families exhibit little overlap in caudal fin PC2 morphospace, with filefishes exhibiting long, low AR caudal fins (high PC2 scores) and triggerfishes almost exclusively exhibiting short, high AR fins (low PC2 scores).

The primary axis of dorsal fin variation (PC1: 69%) differentiates fishes by AR with posteriorly tapering, high AR fins occupying areas of low PC1 morphospace and low AR, more rectangular fins occupying areas of high PC1 morphospace (Spearman’s rank correlation rho = -0.9654, P = 2.20e^-16^) (Figure 5C). Dorsal fin PC2 (13%) primarily describes the angle between the leading edge of the fin and the body axis, with dorsal fins exhibiting anteriorly oriented leading edges possessing high PC2 scores and fins with posteriorly oriented leading edges possessing low PC2 scores. Filefishes cluster in high PC1 morphospace defined by low AR dorsal fins, with the exceptions of *Paramonacanthus sulcatus* and *P. oblongus*, which lie in low PC1 morphospace (high AR fins) primarily occupied by triggerfishes. Both families span the full range of PC2 morphospace.

Anal fin morphospace is similar to that of the dorsal fin, with the primary axis of variation (PC1: 57%) significantly correlated with AR (Spearman’s rank correlation rho = -0.9494, P = 2.20e^-16^) (Figure 5D). High AR fins are associated with low PC1 scores. PC2 (17%) describes the angle of the leading edge of the fin and the distribution of area along the fin, with anteriorly-oriented leading edges and front-loaded area distributions found in high PC2 morphospace, and posteriorly oriented leading edges and back-loaded area distributions found in low PC2 morphospace. Balistoid families show more separation in anal fin morphospace than in dorsal fin morphospace with filefishes almost exclusively exhibiting low AR fins (high PC1) and triggerfishes tending towards mid-to-high AR fins (mid-to-low PC1). Two triggerfish species have diverged from all other balistoids to occupy the top left corner of morphospace (low PC1, high PC2) described by very high AR, sharply posteriorly-tapering anal fins. Interestingly, the filefish *Paramonacanthus oblongus* is the next closest species to this region of morphospace.

### Morphological Integration

Phylogenetic integration tests revealed high levels of morphological integration between all fin and body regions (Table 1, Figure 6) indicating that balistoid fin and body shapes have been tightly correlated throughout their evolutionary history. Pairwise PGLS regressions indicated that dorsal and anal fin shapes are particularly tightly correlated with significant correlations detected between dorsal fin PC1 and anal fin PC1, dorsal fin PC2 and anal fin PC2, and dorsal fin PC2 and anal fin PC1 (P = 2.2e^-16^, multiple R^2^ = 0.6722; P = 0.004353, multiple R^2^ = 0.1047; P = 1.17e^-7^, multiple R^2^ = 0.3173). Additionally, bodies with narrow caudal peduncles and long faces (low body PC2) and truncate-to-forked caudal fins (high caudal fin PC1) are correlated with anal fins that exhibit posteriorly-oriented leading edges and back-loaded area distributions (low anal fin PC2) (P = 8.98e^-5^, multiple R^2^ = 0.1861; P = 0.03859, multiple R^2^ = 0.05731). Finally, high AR dorsal fins with long leading edges (low dorsal fin PC1) are correlated with convex caudal fins (low caudal fin PC1) (P = 0.002223, multiple R^2^ = 0.1211). No significant correlations were found between the primary axes of body and caudal fin shapes or body and dorsal fin shapes.

**Figure 6.**
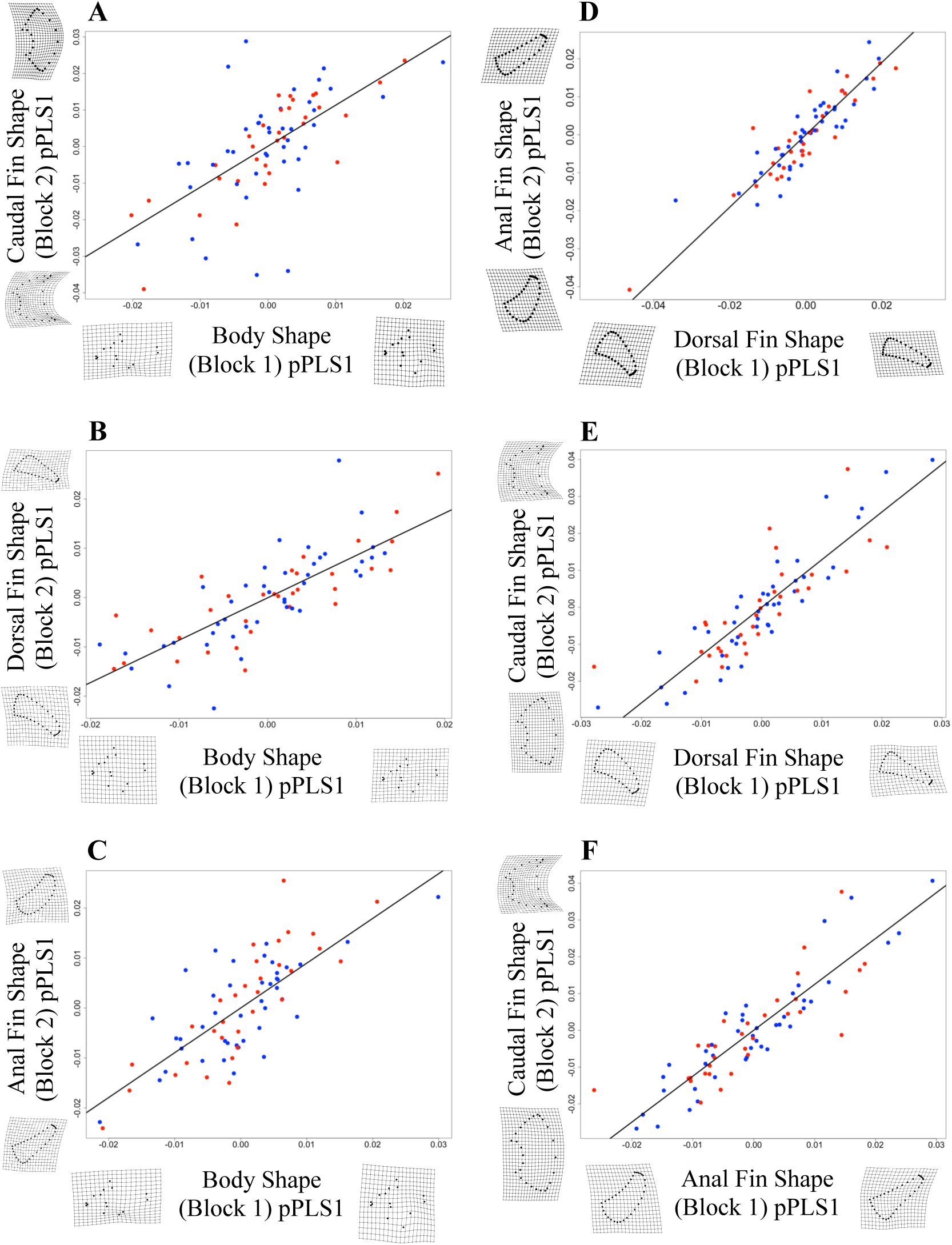
Evolutionary morphological integration. Each panel depicts the relationship between the primary phylogenetic partial least squares (pPLS) axes of shape covariation between pairs of morphological subunits. (A) Body and caudal fin. (B) Body and dorsal fin. (C) Body and anal fin. (D) Dorsal and anal fin. (E) Dorsal and caudal fin. (F) Anal and caudal fin. Points represent species and are colored by family with triggerfishes in red and filefishes in blue.

**Table 1.** Phylogenetic integration and pair-wise correlations between primary shape pPLS axes.

| Shape Dataset<br>(Block 1 vs Block 2) | Overall Phylogenetic<br>Integration (r-PLS/ P value) | Block 1 pPLS1 vs Block 2 pPLS1<br>(Multiple R-squared/ P value) |
| --- | --- | --- |
| Body vs Caudal Fin | 0.6583/ $1.0e^{-4}$ | 0.4334/ $1.8e^{-10}$ |
| Body vs Dorsal Fin | 0.7918/ $1.0e^{-4}$ | 0.6270/ $2.2e^{-16}$ |
| Body vs Anal Fin | 0.7768/ $1.0e^{-4}$ | 0.6034/ $4.1e^{-16}$ |
| Dorsal Fin vs Anal Fin | 0.9012/ $1.0e^{-4}$ | 0.8122/ $2.2e^{-16}$ |
| Dorsal Fin vs Caudal Fin | 0.8682/ $1.0e^{-4}$ | 0.7537/ $2.2e^{-16}$ |
| Anal Fin vs Caudal Fin | 0.8839/ $1.0e^{-4}$ | 0.8839/ $1.0e^{-4}$ |

Legend: Phylogenetic partial least squared is abbreviated pPLS.

### Evolutionary Ecomorphology

Planktivorous balistoid species and species that live in open ocean pelagic habitats exhibit distinct dorsal fin, anal fin, and full shape morphologies compared to the other ecotypes examined (Figures 7 and 8). All morphologies (PC scores and ratio measurements) that distinguished planktivorous and pelagic species from other ecotypes are associated with median fin AR. Open ocean pelagic species exhibit significantly lower full shape PC2 scores than fishes that live on structured coral reefs (t = -4.98, P = 0.006). These pelagic fishes are characterized by high AR fins, short dorsal spines, and shallow ventral keels, while the structured reef fishes possess low AR fins, long dorsal spines, and deep ventral keels (Figure 8).

**Figure 7.**
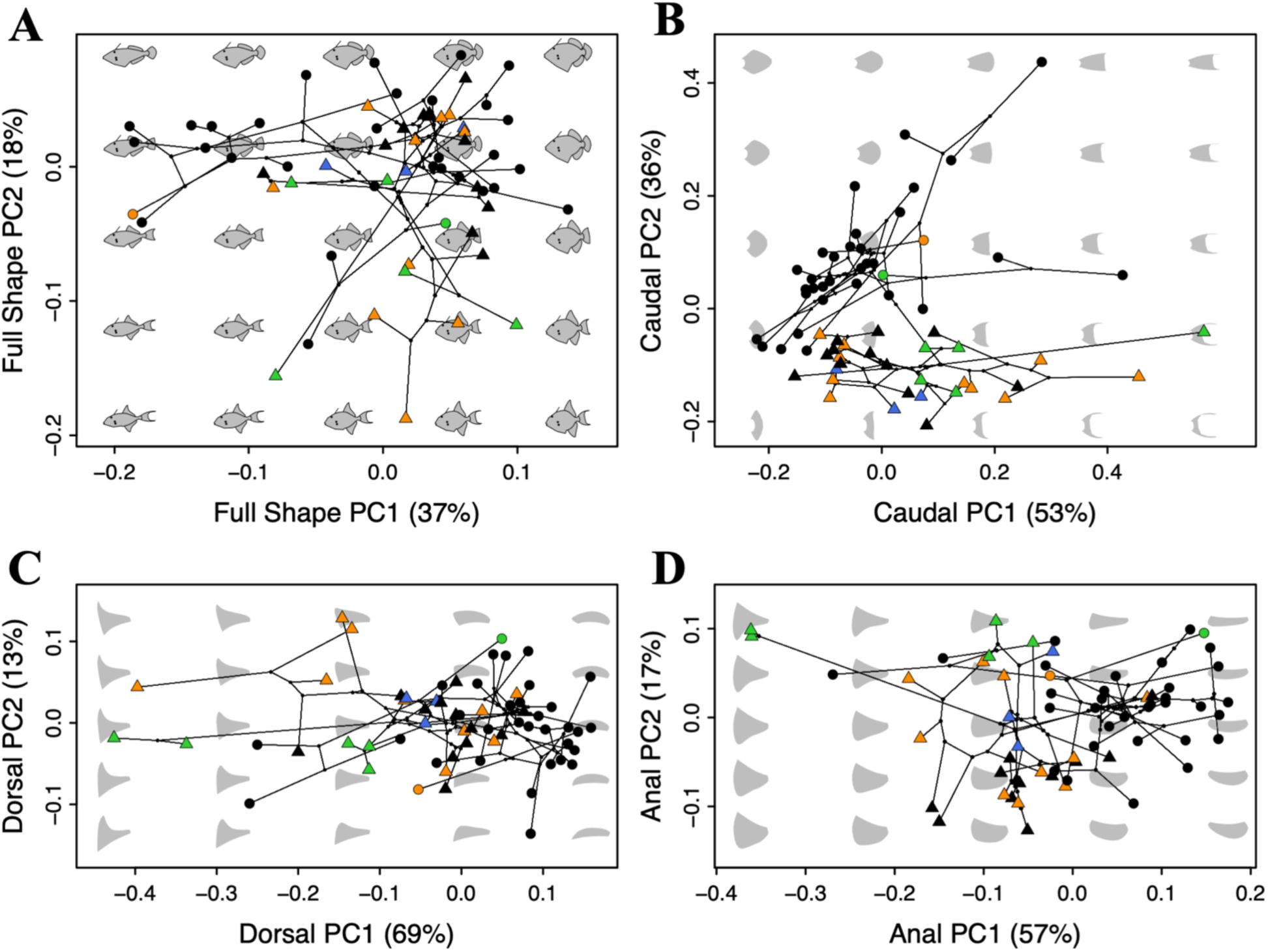
Relationships between balistoid morphology and feeding mode. Backtransformation phylomorphospace colored coded according to the feeding mode of each species with black indicating benthic grazers, orange indicating predators of elusive prey, blue representing mixed grazing and planktivory, and green representing planktivory. Triggerfishes are represented by triangles and filefishes are represented by circles. (A) Full Shape. (B) Caudal Fin. (C) Dorsal Fin. (D) Anal Fin.

**Figure 8.**
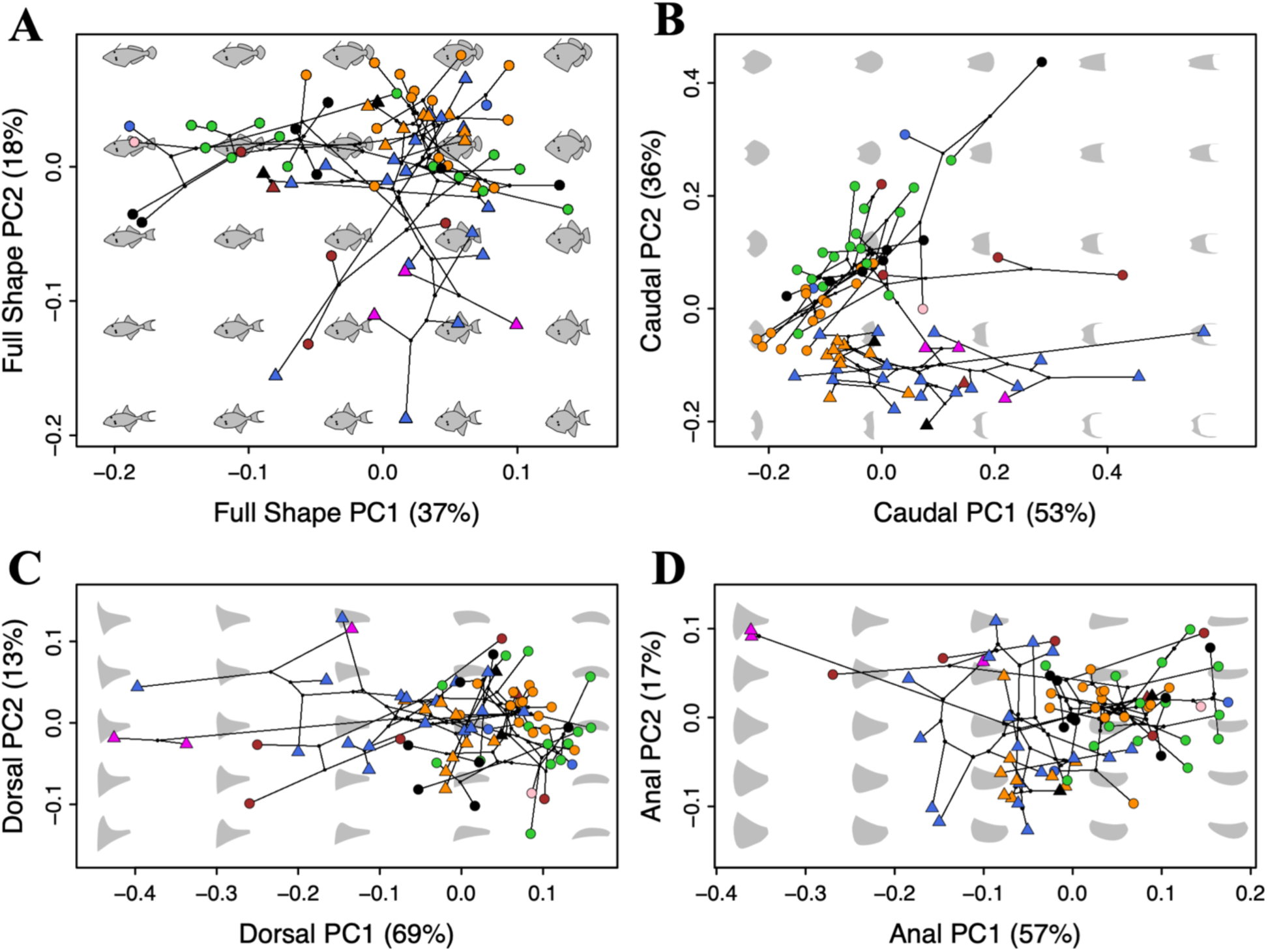
Relationships between balistoid morphology and habitat use. Backtransformation phylomorphospace colored coded according to the primary habitat of each species with orange indicating structured reef, blue indicating open reef, green indicating seagrass/ weedy, black indicating open ocean demersal, brown indicating coastal bare bottom, beige indicating open ocean rafting, and pink indicating open ocean pelagic. Triggerfishes are represented by triangles and filefishes are represented by circles. (A) Full Shape. (B) Caudal Fin. (C) Dorsal Fin. (D) Anal Fin.

Significant differences were found in dorsal fin ARs (log transformed) for both the feeding mode and habitat categories (Table 2). Planktivorous fishes exhibit significantly higher dorsal fin ARs than benthic grazers (p = 0.0066). Likewise, open ocean pelagic fishes possess higher AR dorsal fins than fishes inhabiting the structured reef, open reef, demersal, and seagrass/weedy habitat groups (Table 2). The relationships between dorsal fin AR and ecology are further supported by significant relationships between dorsal fin PC1 (highly correlated with dorsal fin AR) and feeding and habitat groups (Table 2; Figures 7C and 8C). Dorsal fin geometric morphometrics provide more detail about fin shapes associated with these ecotypes than the AR data alone. Specifically, fishes with long leading edges and posteriorly-tapering dorsal fins (low dorsal PC1) are associated with planktivorous and open ocean pelagic habitats.

**Table 2.** Significant results of phylogenetic ANOVA ecomorphology and pairwise t-tests.

| Ecological Variable and Morphological Axis | phlyANOVA Significance (F/ P values) | Primary Ecological Group | Significantly Different Ecological Groups | Pairwise t-test Significance (t/ P value) |
| --- | --- | --- | --- | --- |
| Habitat and log(Full Shape PC2) | 8.276/ 0.0969 | Pelagic | Structured Reef | -4.984/ 0.0060 |
| Feeding Mode and log(Dorsal AR) | 10.0591/ 0.0205 | Planktivory | Benthic Grazer | -4.746/ 0.0066 |
| Habitat and log(Dorsal AR) | 11.766/ 0.0213 | Pelagic | Structured Reef, Open Reef, Seagrass/ Weedy, Demersal | -5.772/ 0.0030;<br>-3.828/ 0.0104;<br>-6.464/ 0.0042;<br>-4.854/ 0.0204 |
| Feeding Mode and Dorsal PC1 | 8.597/ 0.0492 | Planktivory | Benthic Grazer | -4.468 / 0.0180 |
| Habitat and log(Dorsal PC1) | 11.662/ 0.0278 | Pelagic | Structured Reef, Open Reef, Seagrass/ Weedy, Demersal | -5.693/ 0.0015;<br>-3.826/ 0.0117;<br>-6.379/ 0.0070;<br>-4.890/ 0.0168 |
| Feeding Mode and log(Anal Fin AR) | 9.870/ 0.0220 | Planktivory | Benthic Grazer | -5.028/ 0.0036 |
| Habitat and log(Anal Fin AR) | 13.107/ 0.0169 | Pelagic | Structured Reef, Open Reef, Seagrass/ Weedy, Demersal | -6.171/ 0.0015;<br>-4.641/ 0.0026;<br>-7.280/ 0.0015;<br>-5.814/ 0.0036 |
| Habitat and Anal Fin PC1 | 11.985/ 0.0259 | Pelagic | Structured Reef, Open Reef, Seagrass/ Weedy, Demersal | 5.487/ 0.0030;<br>4.333/ 0.0036;<br>6.927/ 0.0030;<br>5.749/ 0.0030 |
| Feeding Mode and Anal Fin PC2 | 6.011/ 0.1793 | Planktivory | Benthic Grazer, Predators of Elusive Prey | -4.106/ 0.0290;<br>-3.751/ 0.0078 |

Phylogenetic ANOVA is abbreviated phlyANOVA. Aspect ratio is abbreviated AR. All P values were adjusted for multiple tests using the Holm method.

Similar trends were found in regard to log-transformed anal fin ARs, with planktivorous fishes and open ocean pelagic fishes exhibiting significantly higher anal fin ARs than nearly all other feeding and habitat groups (Table 2). Anal fin PC1 showed additional between-habitat morphological differences, with open ocean pelagic fishes exhibiting triangular, posteriorly-tapering anal fins (low PC1) and nearly all other habitat groups possessing more rectangular or rounded anal fins (Figure 9D; Table 2). Interestingly, no relationship was discovered between anal fin PC1 and feeding mode, but planktivorous fishes were found to exhibit significantly higher anal fin PC2 scores (posteriorly tapering fins) than benthic grazers and predators of elusive prey (Table 2).

No significant relationships were detected between fin asymmetry and ecology. Furthermore, no significant pair-wise morphological differences were discovered between any other feeding or habitat groups, indicating that benthic grazers, predators of elusive prey, and fishes that combine benthic grazing and planktivory are morphologically indistinguishable from one another, as are fishes that live in structured reef, open reef, seagrass bed, coastal bare bottom, and open ocean demersal habitats.

Hypothesized trends between morphology and ecology were also examined using convergence tests. As predicted, fishes living in structured reef environments were found to converge in morphospace defined by short, deep bodies (high full shape PC1), long dorsal spines and deep ventral keels (high full shape PC2), low dorsal and anal fin ARs (high dorsal and anal fin PC1s), and deep, slightly convex caudal fins (low caudal fin PCs 1 and 2) (Table 3). Pelagic fishes have converged in morphospace defined by high AR, tapering anal fins (high anal fin PCs 1 and 2), high AR dorsal fins (low dorsal fin PC1), and deep, slightly concave caudal fins (low caudal fin PC2 and intermediate PC1). Benthic grazers also exhibited convergence in full, caudal fin, dorsal fin, and anal fin shapes (Table 3), where they generally cluster in areas defined by low AR dorsal and anal fins and convex caudal fins (black shapes in Figure 7). Finally, planktivores have converged in dorsal and anal fin morphospace (Table 3), where they generally exhibit high dorsal fin ARs (high PC1) and elongate, tapering anal fins (high PC2) (Figure 7 C and D).

**Table 3.** Results of ecomorphology convergence tests.

| Test Group (Morphology - Ecology) | C1 | P value |
| --- | --- | --- |
| Full Shape - Offshore Pelagic | 0.220 | 0.06293706 |
| Full Shape - Structured Reef | 0.430 | **1.60e<sup>-7</sup>** |
| Full Shape - Benthic Grazing | 0.219 | **1.60e<sup>-7</sup>** |
| Full Shape - Planktivore | 0.079 | 0.8582834 |
| Dorsal Fin - Offshore Pelagic | 0.427 | **0.047904192** |
| Dorsal Fin - Structured Reef | 0.497 | **1.60e<sup>-7</sup>** |
| Dorsal Fin - Benthic Grazing | 0.297 | **1.60e<sup>-7</sup>** |
| Dorsal Fin - Planktivore | 0.363 | **0.01996008** |
| Anal Fin - Offshore Pelagic | 0.492 | **1.60e<sup>-7</sup>** |
| Anal Fin - Structured Reef | 0.405 | **1.60e<sup>-7</sup>** |
| Anal Fin - Benthic Grazing | 0.214 | **1.60e<sup>-7</sup>** |
| Anal Fin - Planktivore | 0.302 | **0.011976048** |
| Caudal Fin - Offshore Pelagic | 0.747 | **1.60e<sup>-7</sup>** |
| Caudal Fin - Structured Reef | 0.540 | **1.60e<sup>-7</sup>** |
| Caudal Fin - Benthic Grazing | 0.314 | **1.60e<sup>-7</sup>** |
| Caudal Fin - Planktivore | 0.113 | 0.8582834 |

Legend: C1 convergence represents the proportion of the maximum ancestral distance between focal species (species in each ecotype group) that has been closed over evolutionary time. Bold values indicate statistically significant convergence results. All P values were adjusted for multiple tests using the Holm method.

## DISCUSSION

Balistoid fishes exhibit a wide range of body morphologies from deep bodied to elongate forms, possess dorsal and anal fins that vary in aspect ratio by an order of magnitude, and caudal fins that range from shallow and convex to deep and forked. Every morphological unit tested displayed evolutionary integration with every other unit along primary pPLS covariation axes, and the dorsal and anal fins exhibited especially high integration as evidenced by highly correlated primary and secondary PC scores. Despite this strong evolutionary integration, balistoid dorsal and anal fins are asymmetrical in most balistoid fishes, with triggerfishes, on average, displaying higher levels of fin asymmetry than filefishes. Species generally cluster by family in morphospace, however each morphological unit examined (full shape, body, dorsal fin, anal fin, and caudal fin) also exhibits regions of morphospace with substantial overlap of species from both families. Triggerfishes generally have higher AR dorsal, anal, and caudal fins than filefishes, but multiple species from both families exhibit high, medium, and low AR dorsal and anal fins, demonstrating the high level of convergence in these traits.

The diversity of balistoid fin and body shapes is reflected in their habitat and feeding ecologies. Phylogenetic morphometric analysis supports the hypotheses that high AR dorsal and anal fins are associated with planktivorous diets and offshore pelagic habitats. These ecomorphological relationships are congruent with biomechanical predictions from swimming tests in which high AR balistoid fins are shown to be associated with increased endurance swimming performance (George and Westneat 2019). Both planktivory and pelagic habitats present these fishes with the challenge of spending large amounts of time swimming in the open water. Conversely, fishes that live in close association with structured coral reefs and benthic grazing species were found to converge in areas of morphospace defined by low AR fins, characteristics likely to facilitate performance of complex maneuvers beneficial to life in these ecotypes. These strong ecomorphological relationships are concluded to be central drivers of the multiple evolutionary convergence events towards high and low AR dorsal and anal fins in the superfamily Balistoidea. Notably, many species with different habitat and feeding ecologies also inhabit overlapping regions of morphospace, demonstrating that despite some ecomorphological clustering, balistoid fishes are capable of using a wide range of fin and body morphologies to feed on a variety of prey items and inhabit most marine habitats.

### Patterns of Morphological Evolution

Fishes in the superfamily Balistoidea are quite morphologically diverse and exhibit high levels of evolutionary morphological integration among all measured fin and body regions, supporting the previously reported pattern of morphological integration among triggerfishes (Dornburg et al. 2011), and expanding this trend to include filefishes. The full shape dataset revealed that triggerfishes and filefishes appear to have diversified along different major axes of morphological variation (Figure 5A). Filefishes (Monacanthidae) span the full range of length-depth ratios (PC1), and triggerfishes (Balistidae) span nearly the full range of fin aspect ratios (PC2), but exhibit conserved, intermediate length-depth ratios. The families are most dissimilar in caudal fin shape (Figure 5B), with triggerfishes possessing deep, high AR tail fins (low PC2) and filefishes possessing elongate, low AR tail fins (high PC2). Most species in both families possess convex caudal fins (Low PC1), with only a few members of each family possessing truncate or forked tail fins. However, when examining a combined PC1-PC2 caudal fin morphospace, there is virtually no overlap between families. The differences in caudal fin shape appear to be a result of a morphological divergence event very shortly after the Balistidae-Monacanthidae split followed by conserved caudal fin AR evolution within each family. The triggerfish common ancestor likely possessed a high AR caudal fin, as do the majority of extant triggerfishes, with only one species (*Pseudobalistes naufragium*) exhibiting a low AR (Figure 3). In contrast, the filefish common ancestor likely had a low or medium AR caudal fin, and all measured extant filefishes still exhibit one of these two states.

Members of both families overlap considerably in dorsal and anal fin morphospace (Figure 5 C and D), with the vast majority of species possessing low to medium AR fins (mid to high PC1 scores). Although triggerfish species exhibit the highest AR dorsal and anal fins, two filefish species (*Paramonacanthus oblongus* and *Paramonacanthus sulcatus*) have diverged from their low AR relatives to occupy areas of high AR morphospace. Dorsal and anal fin shapes are highly integrated and tend to nearly mirror each other (Figure 6D), yet perfect symmetry in the positioning and shape of these median fins is rare within balistoid fishes (Figure 4). The integration of the dorsal and anal fins is likely a result of reliance on coordinated fin motions between the two fins to power steady balistiform swimming (Dornburg et al. 2011), and this biomechanical coupling makes their asymmetry unusual. Perfect symmetry is quite common in other paired propulsors, such as fish pectoral fins and vertebrate wings, largely because pectoral fins and wings lie across the evolutionarily conserved, bilaterally symmetrical left-right plane of these organisms. Asymmetry of paired locomotor appendages causes uneven thrust generation and shifts in the centers of lift and drag away from the center of mass, resulting in increased roll and yaw around the body axis and reduced turning performance (Thomas 1993; Swaddle and Witter 1998). In fact, bilateral AR symmetry is so conserved in fishes that power locomotion with paired pectoral fins that comparative morphology and performance studies can measure the AR of one pectoral fin and confidently assume an equal AR in the other (Fulton and Bellwood 2004; Denny 2005; Martinez et al. 2016). Likewise, bird and bat wing ARs are typically calculated using a single wing-tip to wing-tip span measurement to encompass both wings, highlighting the well-supported assumption of symmetry in these paired bilateral propulsors (Savile 1957; Norberg and Lighthill 1981).

The paired propulsors utilized by balistoid fishes, on the other hand, lie across the dorsal-ventral axis of the body, an axis much less likely to allow for perfect symmetry in these bilaterally symmetrical organisms. In this system, the anal fin is bounded anteriorly by the position of the anus, while the dorsal fin is free to move as far forward as the back of the head. Consequently, balistoid anal fins tend to possess fewer fin rays, be slightly shorter, and be positioned slightly posteriorly compared to the dorsal fins. Thus, the finding that 40% of the 287 balistoid individuals examined exhibited equal dorsal and anal fin ARs could be regarded as a high degree of symmetry when considered in the broader evolutionary context of dorsal-ventral asymmetry across vertebrate body plans. However, from a functional perspective, this paired-propulsor asymmetry places balistoid fishes in a hydrodynamic situation quite unlike that of the symmetrical pectoral fin and paired flight wing systems. The asymmetry of positioning and shape of most balistoid dorsal and anal fins sets up a biomechanical paired-propulsor system that likely requires slightly different dorsal and anal fin kinematics to avoid constant pitch and roll about the center of mass. Interestingly, no relationships were detected between fin asymmetry and ecology, indicating that fin asymmetry does not appear to hinder or improve balistoid fishes’ abilities to successfully occupy the full range of balistoid habitats and feeding modes. More work on balistoid swimming kinematics is needed to determine how balistoid fishes compensate for this asymmetry.

Ancestral state estimations revealed high levels of convergence in dorsal and anal fin aspect ratios as well as widespread convergence on highly asymmetrical fins, both within and between families. Triggerfishes likely evolved from a common ancestor with moderately asymmetrical, high AR dorsal and anal fins and exhibit many independent convergence events towards medium and low AR fins, while filefishes likely evolved from a common ancestor with fairly symmetrical, low AR fins and exhibit multiple convergence events onto medium and high AR fins. These morphological transitions are likely accompanied by changes in fin kinematics during steady swimming, as balistoid fishes are known to lie on a kinematic continuum from highly undulatory, wave-like dorsal and anal fin kinematics in low AR fins to highly oscillatory, flapping dorsal and anal fin kinematics in high AR fins (Wright 2000). Convergent evolution towards highly asymmetrical fins in fishes possessing both high and low AR fins suggest that balistoid species are able to overcome instabilities introduced by fin asymmetries regardless of whether they use undulatory or oscillatory fin kinematic strategies. The combination of fin shape and fin kinematics is also likely to affect the maneuverability of balistoid fishes, with low-AR, undulatory fins providing a strong maneuverability advantage over high-AR, flapping fins.

### Evolutionary Ecomorphology

Many past studies have hypothesized ecomorphological relationships between balistoid fin shapes and ecology (Wright 2000; Dornburg et al. 2011; George and Westneat 2019), largely based on relationships between fin aspect ratios and swimming performance. In this study, two distinct comparative methods revealed many interesting ecomorphological relationships among balistoid fishes. Phylogenetic ANOVAs revealed that offshore-pelagic and planktivorous fishes lie in fairly exclusive areas of morphospace defined by higher AR and more posteriorly-tapering dorsal and anal fins than species with other habitat and feeding mode ecologies (Figures 7 and 8). These relationships between high AR fins and pelagic and planktivorous fishes are supported by the strong biomechanical correlation between high AR fins and increased balistoid endurance swimming performance required for successful occupation of these ecotypes. The only other species to occupy this area of morphospace occur in either open reef or coastal bare bottom habitats (Figure 8), both open environments that would benefit from high endurance swimming performance. Interestingly, open reef and coastal bare bottom species also occupy nearly all other areas of morphospace, indicating that high AR fins may be beneficial, but are not necessary for life in these environments.

We predicted that fishes in the structured reef habitat group and the benthic grazing feeding group would exhibit lower AR median fins, longer dorsal spines, and deeper ventral keels than fishes in other ecotypes, but these hypotheses were not supported by the phylogenetic ANOVA tests. These morphological characteristics were in fact shared by species across nearly all balistoid feeding ecologies and habitat types. However, we used convergence tests to identify morphological clusters of species from a single ecotype rather than testing whether these species have evolved to occupy exclusive areas of morphospace. These tests revealed that benthic grazing species have indeed converged on low AR dorsal and anal fins and elongate, convex caudal fins of intermediate-to-low AR (Figure 7), features likely to facilitate high maneuverability and smooth backwards swimming beneficial for grazing on irregularly shaped coral and rocks. Additionally, fishes living in structured reef environments have converged on long dorsal spines (high full shape PC2) and deep ventral keels (high full shape PC1), two mobile elements that allow these fishes to safely lock themselves into cervices in their structurally complex habitats (Figure 8). Structured reef-associated fishes have also converged on maneuverable, low AR dorsal and anal fins and high burst performance deep caudal fins. Similar trends have been reported in wrasses, in which low AR pectoral fins are associated with sheltered reef areas and high AR fins are associated with high wave energy areas (Fulton et al. 2001).

At least one member from every feeding and habitat group except the offshore pelagic group possesses low AR dorsal and anal fins, including one planktivorous filefish *Arotrolepis filicauda* (Figures 7 and 8). This extensive morphological convergence towards low AR dorsal and anal fins across ecotypes suggests that low AR fins are highly versatile in a number of ecological contexts. Many balistoid species exhibit mating and parental care behaviors that involve a high degree of maneuverability such as swimming in tight circles with their competitors (Kawase and Nakazono 1996), constructing nests (Fricke 1980; Gladstone 1994), and guarding and tending to benthic eggs (Fricke 1980; Nakazono and Kawase 1993; Kawase and Nakazono 1995; Ishihara and Kuwamura 1996; Kawase 2003), behaviors likely requiring the maneuverability conferred by low AR fins. Numerous open reef and planktivorous fishes display some form of these mating and parental care behaviors, which may explain the presence of low AR fins in these ecotypes despite their extensive time spent in the water column. Future research on relationships between morphology and mating and parental care behaviors may help elucidate the high number of convergence events on low AR dorsal and anal fins across ecotypes.

## ACKNOWLEDGEMENTS

The following museums and databases provided access to specimens or photographs used in morphometric analyses: Kanagawa Prefectural Museum of Natural History (KPM), Field Museum of Natural History (FMNH), University of Michigan Museum of Zoology (UMMZ), Bernice Pauahi Bishop Museum (BPBM), US National Museum of Natural History (NMNH), French National Museum of Natural History (MNHN), The Australian Museum (AM), Florida Museum of Natural History (FLMNH), Digital Fish Library at the University of California San Diego (DFL), Western Australian Museum (WAM), Harvard Museum of Comparative Zoology (MCZ), Academy of Natural Sciences (ANSP), California Academy of Sciences (CAS), Bold Systems, GBIF, FishBase, and Royal Ontario Museum (ROM). We thank Hiroaki Hayashi and Hiroshi Senou of the KPM for measuring and photographing specimens. We thank Douglas Nelson (UMMZ) and Caleb McMahan, Susan Mochel, and Kevin Swagel (FMNH) for providing lab space and specimens. We thank Nicholas Slimmon for help digitizing photographs, Aaron Olsen, Meg Malone, and Isaac Krone for morphometrics advice, and Melina Hale, Callum Ross, Chloe Nash, Lily Hughes, Katie Whitlow, and Sam Gartner for advice on the writing of this manuscript.

