## Supplemental Methods for "Fin Shape, Asymmetry, and Evolutionary Ecomorphology in Triggerfishes and Filefishes (Superfamily: Balistoidea)"

**Habitat Use and Feeding Traits.** Habitat category assignments were based on descriptive data and/or catch records, with quantitative data (timed behavioral studies) available for a few species. Quantitative diet data (based on stomach contents or timed feeding behavior studies) were available for 46 of the 80 species, allowing for easy placement of these species into primary feeding groups. When quantitative diet data was not available (n = 34 species), feeding mode classifications were based on detailed published descriptions of feeding behaviors, usually confirmed by multiple sources for each species (Table S1). Habitat classifications were made for all 80 balistoid species in this study, and feeding mode classifications were made for 70 species.

**Morphometrics.** All specimens included in this study were scaled (using a ruler in the photograph or specimen-specific length information) with the exception of 7 visibly adult specimens due to the lack of accessible, scaled specimens for these species (see Table S2). For the 7 cases in which specimen-specific scale data were not available, and considering that shape and not size scaling is explored here, each specimen was assigned a common adult standard length reported for their species on *FishBase* (Froese and Pauly 2019).

**Integration**: Morphological PLS analysis differs from morphological PCA in that the primary PLS axes correspond to the axes of maximum morphological *covariation* between two selected morphological units (such as dorsal and anal fins), termed blocks, while the primary PCA axes correspond to the axes encompassing the maximum morphological variance between species within a single morphological unit (ie. dorsal fin only). Thus, primary PLS axes represent the axis of morphological variation in one morphological subunit (ie. the dorsal fin) that is mostcorrelated with variation in the other morphological subunit (ie. the anal fin). This makes PLS analysis a powerful tool for identifying significant integration between morphological subunits along *any* axes of variation, while PGLS correlations between the primary or secondary PC axes (PCs 1 and 2) of two morphological units goes one step further by demonstrating that the two subunits display evolutionary integration along their *most significant* axes of morphological variation.

**Statistical Analysis Notes.** All statistical analyses were carried out in R version 4.0.2 (R Core Team 2020). Prior to running all statistical tests described above, data and residuals were examined for alignment with the assumptions of each test. Phylogenetic generalized least squares (PGLS) and phylogenetic ANOVA analyses both assume that the residuals of the analyses are normally distributed. This assumption was assessed by visualizing the distribution of residuals from each test and performing Lilliefors (Kolmogorov-Smirnov) tests using the *lillie.test* function in the *nortest* R package (Gross and Ligges 2015). The phylogenetic ANOVA tests also assume homogeneity of variance (equal variance across groups), which was tested using Levene's tests with the *leveneTest* function in the *car* R package (Fox and Weisberg 2011). Log-transformations were used to improve adherence to these assumptions when assumptions were violated using raw data.

R Core Team. 2020. R: A Language and Environment for Statistical Computing. R Foundation for Statistical Computing, Vienna, Austria.
