## Supplemental Table 1 for "Fin Shape, Asymmetry, and Evolutionary Ecomorphology in Triggerfishes and Filefishes (Superfamily: Balistoidea)"

Table S1. Balistoid species habitat and feeding ecology classifications and citations.

| Species | Family | Habitat | Habitat Citation | Feeding Mode | Feeding Mode Citation |
|---|---|---|---|---|---|
| *Abalistes stellaris* | Balistidae | Coastal Bare Bottom | (K Matsuura 2001) | BG & EP * | (Kulbicki et al. 2005) |
| *Abalistes stellatus* | Balistidae | Open Ocean Demersal | (Khalaf and Zajonz 2007; Kuiter 2005) | BG | (Carpenter et al. 1997) |
| *Acanthaluteres spilomelanurus* | Monacanthidae | Seagrass/ Weedy | (Scott 1981) | BG * | (Scott 1981) |
| *Acanthaluteres vittiger* | Monacanthidae | Seagrass/ Weedy | (Edgar, Barrett, and Morton 2004; Hyndes et al. 2003) | BG * | (Jones 1992; Last 1975) |
| *Acreichthys tomentosus* | Monacanthidae | Seagrass/ Weedy | (Nakamura and Sano 2004; Peristiwady and Geistdoerfer 1991) | BG * | (Horinouchi et al. 2012; Nakamura et al. 2003; Peristiwady and Geistdoerfer 1991) |
| *Aluterus heudelotii* | Monacanthidae | Open Ocean Demersal | (Wenner 1983) | BG | (K Matsuura 2001) |
| *Aluterus monoceros* | Monacanthidae | Open Ocean Rafting | (Kuiter 1992; Lieske and Myers 2002) | BG * | (Ghosh et al. 2011; Lopez-Peralta and Arcila 2002) |
| *Aluterus schoepfii* | Monacanthidae | Seagrass/ Weedy | (Lieske and Myers 2002; Randall 1967) | BG * | (K Matsuura 2001; Randall 1967) |
| *Aluterus scriptus* | Monacanthidae | Open Reef | (Dominici-Arosemena and Wolff 2006; Lieske and Myers 2002; Randall 1967) | BG * | (Randall 1967; Randall, Allen, and Steene 1990; Wulff, 1994) |
| *Amanses scopas* | Monacanthidae | Structured Reef | (Kuiter 1992; Myers 1991) | BG | (Durville, Chabanet, and Quod 2003; Schroeder 1980) |
| *Arotrolepis filicauda* | Monacanthidae | Coastal Bare Bottom | (Blaber, Brewer, and Harris 1994) | PL * | (Bulman et al. 2001) |

| | | | | | |
|---|---|---|---|---|---|
| *Balistapus undulatus* | Balistidae | Structured Reef | (Hiatt and Strasburg 1960; Lieske and Myers 2002; McClanahan and Shafir 1990; Randall, Allen, and Steene 1990) | BG & EP * | (Hiatt and Strasburg 1960; Randall 1955; Randall, Allen, and Steene 1990) |
| *Balistes capriscus* | Balistidae | Open Ocean Pelagic | (Longley 1941) | BG & EP * | (Goldman, Glasgow, and Falk 2016; Lieske and Myers 2002; Vose and Nelson 1994) |
| *Balistes polylepis* | Balistidae | Open Reef | (Burgess et al. 1984; Dominici-Arosemena and Wolff 2006) | BG & EP * | (Abitia Cárdenas, Rodríguez Romero, and Galván Magaña 1990; Glynn, Stewart, and McCosker 1972) |
| *Balistes punctatus* | Balistidae | Open Reef | (Froese and Pauly 2019) | BG & EP * | (Aggrey-Fynn 2007) |
| *Balistes vetula* | Balistidae | Open Reef | (Hobson 1965; Randall 1967) | BG & EP * | (Hobson 1965; Randall 1967; Turingan 1994) |
| *Balistoides conspicillum* | Balistidae | Open Reef | (Kuiter 1992; Randall, Allen, and Steene 1990) | BG | (Meyer 1985; Schroeder 1980) |
| *Balistoides viridescens* | Balistidae | Structured Reef | (Hiatt and Strasburg 1960) | BG | (Hiatt and Strasburg 1960; Myers 1991; Randall 1980) |
| *Brachaluteres jacksonianus* | Monacanthidae | Seagrass/ Weedy | (Hyndes et al. 2003; Kawase 2005) | BG * | (Last 1975; Scott 1981) |

| | | | | | |
|---|---|---|---|---|---|
| *Cantherhines dumerilii* | Monacanthidae | Open Reef | (Hobson 1974; Myers 1991) | BG * | (Disalvo, Randall, and Cea 2007; Hiatt and Strasburg 1960; Hobson 1974) |
| *Cantherhines fronticinctus* | Monacanthidae | Structured Reef | (Hutchins and Randall 1982; Lieske and Myers 2002; Taquet et al. 2017) | BG | (Taquet et al. 2017) |
| *Cantherhines macrocerus* | Monacanthidae | Structured Reef | (Lieske and Myers 2002; Randall 1964; 1967) | BG * | (Randall and Hartman 1968; Randall 1967; Turingan, Wainwright, and Hensley 1995; Wulff, 1994) |
| *Cantherhines pardalis* | Monacanthidae | Structured Reef | (Schroeder 1980; Taquet et al. 2017) | BG | (Kawase and Nakazono 1994) |
| *Cantherhines pullus* | Monacanthidae | Structured Reef | (Lieske and Myers 2002; Randall 1967) | BG * | (Randall and Hartman 1968; Randall 1967) |
| *Cantherhines sandwichiensis* | Monacanthidae | Structured Reef | (Hobson 1974) | BG * | (Hobson 1974; Randall 1985) |
| *Cantherhines verecundus* | Monacanthidae | Structured Reef | (Randall 2007) | BG * | (Hutchins and Randall 1982) |
| *Canthidermis maculata* | Balistidae | Open Ocean Pelagic | (Berry and Baldwin 1966; Lieske and Myers 2002; Taquet et al. 2007) | PL | (Bacchet, Zysman, and Lefèvre 2016; Lieske and Myers 2002; Sandin and Williams 2010) |
| *Canthidermis sufflamen* | Balistidae | Open Ocean Pelagic | (Brito, Falcon, and Herrera 1995; Myers 1991) | PL * | (Randall 1967) |
| *Chaetodermis penicilligerus* | Monacanthidae | Seagrass/ Weedy | (Kuiter 1992; Lieske and Myers 2002; Schroeder 1980) | BG | (Schroeder 1980) |
| *Eubalichthys mosaicus* | Monacanthidae | Seagrass/ Weedy | (Taquet et al. 2017) | BG | (Taquet et al. 2017) |

| | | | | | |
|---|---|---|---|---|---|
| *Melichthys indicus* | Balistidae | Open Reef | (Randall 1995) | PL & BG | (Patankar et al. 2018; Randall 1995) |
| *Melichthys niger* | Balistidae | Open Reef | (Hobson 1974; Randall 1955; Sancho, Petersen, and Lobel 2000) | PL & BG * | (Hobson 1974; Randall 1967)h |
| *Melichthys vidua* | Balistidae | Structured Reef | (Hiatt and Strasburg 1960; Sancho, Petersen, and Lobel 2000) | BG & EP | (Hiatt and Strasburg 1960; Lieske and Myers 2002; Randall, Allen, and Steene 1990) |
| *Meuschenia freycineti* | Monacanthidae | Seagrass/ Weedy | (Baker 2011; Edgar, Barrett, and Morton 2004; Valesini, Potter, and Clarke 2004) | BG * | (Bell, Burchmore, and Pollard 1978; Bulman et al. 2001; Burchmore, Pollard, and Bell 1984; Last 1975) |
| *Meuschenia hippocrepis* | Monacanthidae | Seagrass/ Weedy | (Harvey et al. 2012; Taquet et al. 2017) | BG * | (Rodgers, Linnane, and Huveneers 2013) |
| *Meuschenia trachylepis* | Monacanthidae | Seagrass/ Weedy | (Conacher, Lanzing, and Larkum 1979; Wressnig and Booth 2008) | BG * | (Bell, Burchmore, and Pollard 1978) |
| *Monacanthus chinensis* | Monacanthidae | Seagrass/ Weedy | (Bell, Burchmore, and Pollard 1978; Conacher, Lanzing, and Larkum 1979) | BG * | (Bell, Burchmore, and Pollard 1978; Burchmore, Pollard, and Bell 1984; Conacher, Lanzing, and Larkum 1979) |

| | | | | | |
|---|---|---|---|---|---|
| *Monacanthus ciliatus* | Monacanthidae | Seagrass/ Weedy | (Lieske and Myers 2002; Randall 1967; Tabb and Manning 1961) | BG * | (Clements and Livingston 1983; Randall 1967; Springer and Woodburn 1960) |
| *Monacanthus tuckeri* | Monacanthidae | Seagrass/ Weedy | (Layman and Silliman 2002; Lieske and Myers 2002) | BG * | (Lieske and Myers 2002; Randall 1967) |
| *Nelusetta ayraud* | Monacanthidae | Open Ocean Demersal | (Lindholm 1984) | BG & EP * | (Burchmore, Pollard, and Bell 1984; Lindholm 1984) |
| *Odonus niger* | Balistidae | Open Reef | (Fricke 1980; Lieske and Myers 2002; Randall, Allen, and Steene 1990) | PL | (Fricke 1980; Randall, Allen, and Steene 1990) |
| *Oxymonacanthus longirostris* | Monacanthidae | Structured Reef | (Hiatt and Strasburg 1960; Randall 1955) | BG * | (Hiatt and Strasburg 1960; Myers 1991; Sano, Shimizu, and Nose 1984) |
| *Paraluteres prionurus* | Monacanthidae | Structured Reef | (Kuiter 2005; Myers 1991) | BG | (Froese and Pauly 2019; Taquet et al. 2017) |
| *Paramonacanthus choirocephalus* | Monacanthidae | Coastal Bare Bottom | (Hutchins 1997; Travers et al. 2010) | BG | (Carpenter et al. 1997) |
| *Paramonacanthus oblongus* | Monacanthidae | Coastal Bare Bottom | (Carpenter et al. 1997) | BG | (Carpenter et al. 1997) |
| *Paramonacanthus sulcatus* | Monacanthidae | Coastal Bare Bottom | (Froese and Pauly 2019) | BG | (Hutchins 1997) |
| *Pervagor aspricaudus* | Monacanthidae | Structured Reef | (Kuiter 2005; Lieske and Myers 2002; Randall 2007; Taquet et al. 2017) | NA | |
| *Pervagor janthinosoma* | Monacanthidae | Structured Reef | (Myers 1991; Taquet et al. 2017) | NA | |

| | | | | | |
|---|---|---|---|---|---|
| *Pervagor melanocephalus* | Monacanthidae | Structured Reef | (Kuiter 1992; Lieske and Myers 2002) | NA | |
| *Pervagor nigrolineatus* | Monacanthidae | Structured Reef | (Lieske and Myers 2002) | NA | |
| *Pervagor spilosoma* | Monacanthidae | Structured Reef | (Hobson 1974; Lieske and Myers 2002) | BG * | (Hobson 1974; Randall 1985) |
| *Pseudobalistes flavimarginatus* | Balistidae | Open Reef | (Hiatt and Strasburg 1960; Randall, Allen, and Steene 1990) | BG * | (Hiatt and Strasburg 1960; Sano, Shimizu, and Nose 1984) |
| *Pseudobalistes fuscus* | Balistidae | Open Reef | (Hiatt and Strasburg 1960; Myers 1991) | BG * | (Hiatt and Strasburg 1960; Kulbicki et al. 2005; Taquet et al. 2017) |
| *Pseudobalistes naufragium* | Balistidae | Open Reef | (Dominici-Arosemena and Wolff 2006; Witman, Smith, and Novak 2017) | BG * | (Guzmán 1988; Witman, Smith, and Novak 2017) |
| *Pseudomonacanthus macrurus* | Monacanthidae | Seagrass/ Weedy | (Kuiter 1992; Lieske and Myers 2002) | NA | |
| *Pseudomonacanthus peroni* | Monacanthidae | Coastal Bare Bottom | (Blaber, Brewer, and Harris 1994) | NA | |
| *Rhinecanthus abyssus* | Balistidae | Open Ocean Demersal | (Keiichi Matsuura and Shiobara 1989) | NA | |
| *Rhinecanthus aculeatus* | Balistidae | Open Reef | (Hiatt and Strasburg 1960; Lecchini and Galzin 2005; Randall 1955) | BG & EP * | (Hiatt and Strasburg 1960; Lieske and Myers 2002; Myers 1991; Sano, Shimizu, and Nose 1984) |
| *Rhinecanthus assasi* | Balistidae | Open Reef | (Lieske and Myers 2002; Taquet et al. 2017) | BG | (Froese and Pauly 2019) |

| | | | | | |
|---|---|---|---|---|---|
| *Rhinecanthus lunula* | Balistidae | Structured Reef | (Lieske and Myers 2002; Randall, Allen, and Steene 1990) | BG & EP | (Bacchet, Zysman, and Lefèvre 2016) |
| *Rhinecanthus rectangulus* | Balistidae | Structured Reef | (Hiatt and Strasburg 1960; Hobson 1974) | BG * | (Hiatt and Strasburg 1960; Hobson 1974; Randall 1985; Sano, Shimizu, and Nose 1984) |
| *Rhinecanthus verrucosus* | Balistidae | Structured Reef | (K Matsuura 2001; Randall, Allen, and Steene 1990) | BG | (Schroeder 1980) |
| *Rudarius ercodes* | Monacanthidae | Seagrass/ Weedy | (Horinouchi et al. 2013; Kikuchi 1974) | BG * | (Horinouchi et al. 1998; Kawase and Nakazono 1995; Kwak, Huh, and Choi 2006) |
| *Scobinichthys granulatus* | Monacanthidae | Seagrass/ Weedy | (Hyndes et al. 2003; T. Smith, Jenkins, and Hutchinson 2012) | BG * | (Brown, Lewis, and Baker 2008; Burchmore, Pollard, and Bell 1984) |
| *Stephanolepis aurata* | Monacanthidae | Open Ocean Demersal | (Froese and Pauly 2019) | NA | |
| *Stephanolepis cirrhifer* | Monacanthidae | Seagrass/ Weedy | (Kikuchi 1974)K | BG * | (Kawase and Nakazono 1996; Kikuchi 1974; Kwak, Baeck, and Huh 2003) |
| *Stephanolepis hispidus* | Monacanthidae | Open Ocean Demersal | (Longley 1941; Wenner 1983) | BG * | (Livingston 1982; Mancera-Rodríguez and Castro-Hernández 2015) |
| *Stephanolepis setifer* | Monacanthidae | Seagrass/ Weedy | (Lieske and Myers 2002; K Matsuura 2001) | BG | (K Matsuura 2001) |

| | | | | | |
|---|---|---|---|---|---|
| *Sufflamen albicaudatum* | Balistidae | Open Reef | (Lieske and Myers 2002; Taquet et al. 2017) | NA | |
| *Sufflamen bursa* | Balistidae | Structured Reef | (Hobson 1974; Lieske and Myers 2002; Myers 1991) | BG * | (Hobson 1974; Lieske and Myers 2002; Randall 1985) |
| *Sufflamen chrysopterum* | Balistidae | Structured Reef | (Nakamura and Sano 2004; Randall, Allen, and Steene 1990; Taquet et al. 2017) | BG | (Myers 1991; Patankar et al. 2018) |
| *Sufflamen fraenatum* | Balistidae | Open Reef | (Lieske and Myers 2002; Myers 1991; Randall et al. 1993) | BG & EP * | (John E. Randall, Earle, Pyle, Parrish, & Hayes, 1993) |
| *Sufflamen verres* | Balistidae | Structured Reef | (Dominici-Arosemena and Wolff 2006; Hobson 1965) | BG * | (Dominici-Arosemena and Wolff 2006; Glynn, Stewart, and McCosker 1972; Hobson 1965) |
| *Thamnaconus arenaceus* | Monacanthidae | Open Ocean Demersal | (Froese and Pauly 2019) | NA | |
| *Thamnaconus fajardoi* | Monacanthidae | Open Ocean Demersal | (J. L. B. Smith, Smith, and Heemstra 1986) | NA | |
| *Thamnaconus modestoides* | Monacanthidae | Open Ocean Demersal | (Carpenter et al. 1997; K Matsuura and Tyler 1997) | NA | |
| *Thamnaconus modestus* | Monacanthidae | Seagrass/ Weedy | (Masuda, Yamashita, and Matsuyama 2008) | BG * | (Kim, Choi, and Park 2013) |
| *Thamnaconus tessellatus* | Monacanthidae | Open Ocean Demersal | (K Matsuura and Tyler 1997) | NA | |

| | | | | | |
|---|---|---|---|---|---|
| *Xanthichthys auromarginatus* | Balistidae | Open Reef | (K Matsuura 2001; Myers 1991; Taquet et al. 2017) | PL * | (Myers 1991; Randall, Matsuura, and Zama 1978; Randall 2007) |
| *Xanthichthys mento* | Balistidae | Open Reef | (Kawase 2003; Lieske and Myers 2002) | PL * | (Disalvo, Randall, and Cea 2007; Kawase 2003; Randall 1985) |
| *Xanthichthys ringens* | Balistidae | Open Reef | (Lieske and Myers 2002; Randall, Matsuura, and Zama 1978) | PL & BG * | (Turingan, Wainwright, and Hensley 1995) |

Key: BG = Benthic Grazer; EP = Elusive Prey; PL = Planktivore; * = Quantitative data

Bacchet, Philippe, Thierry Zysman, and Yves Lefèvre. 2016. *Guide des poissons de Tahiti et ses îles*. 4th ed. Tahiti, Polynésie Francaise: Au Vent des Îles.

Baker, Janine. 2011. *Reef Fishes of Conservation Concern in South Australia - A Field Guide.* Booklet produced with support from the Adelaide and Mt Lofty Ranges Natural Resources Management Board, South Australia.

Bell, Johann, JJ Burchmore, and David Pollard. 1978. "Feeding Ecology of Three Sympatric Species of Leatherjackets (Pisces : Monacanthidae) from a Posidonia Seagrass Habitat in New South Wales." *Marine and Freshwater Research* 29 (January): 631–43. https://doi.org/10.1071/MF9780631.

Berry, Frederick H., and Wayne J. Baldwin. 1966. "Triggerfishes (Balistidae) of the Eastern Pacific." *Proceedings of the California Academy of Sciences, 4th Series* 34: 429–74.

Blaber, S. J. M., D. T. Brewer, and A. N. Harris. 1994. "Distribution, Biomass and Community Structure of Demersal Fishes of the Gulf of Carpentaria, Australia." *Australian Journal of Marine and Freshwater Research* 45 (3): 375–96. https://doi.org/10.1071/MF9940375.

Brito, A., J. M. Falcon, and R. Herrera. 1995. "Occurrence of the Ocean Triggerfish in the Eastern Atlantic (Canary Islands)." *Journal of Fish Biology* 47 (6): 1099–1101. https://doi.org/10.1111/j.1095-8649.1995.tb06033.x.

Brown, A., J. Lewis, and Janine Baker. 2008. "Uncommon and Cryptic Reef Fishes: Results of Pilot Surveys along Fleurieu Peninsula."

https://www.academia.edu/3718393/UNCOMMON_and_CRYPTIC_REEF_FISHES_RESULTS_OF_PILOT_SURVEYS_ALONG_FLEURIEU_PENINSULA.

Bulman, C., F. Althaus, X. He, N. J. Bax, and A. Williams. 2001. “Diets and Trophic Guilds of Demersal Fishes of the South-Eastern Australian Shelf.” *Marine and Freshwater Research* 52 (4): 537–48. https://doi.org/10.1071/mf99152.

Burchmore, J. J., D. A. Pollard, and J. D. Bell. 1984. “Community Structure and Trophic Relationships of the Fish Fauna of an Estuarine Posidonia Australis Seagrass Habitat in Port Hacking, New South Wales.” *Aquatic Botany* 18 (1): 71–87. https://doi.org/10.1016/0304-3770(84)90081-0.

Burgess, Warren E., Herbert R. Axelrod, Gerald R. Allen, and Roger C. Steene. 1984. *Pacific Marine Fishes. 8: Fishes of California and Western Mexico*. Neptune City, N. J: T. F. H. Publ.

Carpenter, Kent E., Friedhelm Krupp, David Jones, and Uwe Zajonz, eds. 1997. *Living Marine Resources of Kuwait, Eastern Saudi Arabia, Bahrain, Qatar, and the United Arab Emirates*. FAO Species Identification Field Guide for Fishery Purposes. Rome: Food and Agriculture Organization of the United Nations.

Clements, William H., and Robert J. Livingston. 1983. “Overlap and Pollution-Induced Variability in the Feeding Habits of Filefish (Pisces: Monacanthidae) from Apalachee Bay, Florida.” *Copeia* 1983 (2): 331–38. https://doi.org/10.2307/1444375.

Conacher, M. J., W. J. R. Lanzing, and A. W. D. Larkum. 1979. “Ecology of Botany Bay. II. Aspects of the Feeding Ecology of the Fanbellied Leatherjacket, Monacanthus Chinensis (Pisces : Monacanthidae), in Pisidonia Australis Seagrass Beds in Quibray Bay, Botany Bay, New South Wales.” *Marine and Freshwater Research* 30 (3): 387–400. https://doi.org/10.1071/mf9790387.

Disalvo, L. H., John E. Randall, and A. Cea. 2007. “Stomach Contents and Feeding Observations of Some Easter Island Fishes.” *Atoll Research Bulletin* 548: 1–22.

Dominici-Arosemena, Arturo, and Matthias Wolff. 2006. “Reef Fish Community Structure in the Tropical Eastern Pacific (Panamá): Living on a Relatively Stable Rocky Reef Environment.” *Helgoland Marine Research* 60 (4): 287–305. https://doi.org/10.1007/s10152-006-0045-4.

Durville, P, Pascale Chabanet, and Jean-Pascal Quod. 2003. “Visual Census of the Reef Fishes in the Natural Reserve of the Glorieuses Islands (Western Indian Ocean).” *Western Indian Ocean J. Mar. Sci.* 2 (January): 95–104.

Edgar, Graham J., Neville S. Barrett, and Alastair J. Morton. 2004. “Patterns of Fish Movement on Eastern Tasmanian Rocky Reefs.” *Environmental Biology of Fishes* 70 (3): 273–84. https://doi.org/10.1023/B:EBFI.0000033342.89719.39.

Fricke, Hans W. 1980. “Mating Systems, Maternal and Biparental Care in Triggerfish (Balistidae).” *Zeitschrift Für Tierpsychologie* 53 (2): 105–22. https://doi.org/10.1111/j.1439-0310.1980.tb01043.x.

Froese, R, and D Pauly. 2019. “FishBase.” World Wide Web electronic publication. FishBase. 2019. www.fishbase.org.

Ghosh, Shubhadeep, R Thangavelu, Gulshad Mohammed, H Dhokia, Mahendrasinh Zala, Y Savaria, J Polara, and A Ladani. 2011. “Sudden Emergence of Fishery and Some Aspects of Biology and Population Dynamics of Aluterus Monoceros (Linnaeus, 1758) at Veraval." *Indian Journal of Fisheries* 58 (1): 31–34.

Glynn, Peter W., Robert H. Stewart, and John E. McCosker. 1972. "Pacific Coral Reefs of Panamá: Structure, Distribution and Predators." *Geologische Rundschau* 61 (2): 483–519. https://doi.org/10.1007/BF01896330.

Goldman, Sarah F., Dawn M. Glasgow, and Michelle M. Falk. 2016. "Feeding Habits of 2 Reef-Associated Fishes, Red Porgy (Pagrus Pagrus) and Gray Triggerfish (Balistes Capriscus), off the Southeastern United States." *Fishery Bulletin* 114 (3): 317–29. https://doi.org/10.7755/FB.114.3.5.

Guzmán, Héctor M. 1988. "Distribución y abundancia de organismos coralívoros en los arrecifes coralinos de la Isla del Caño, Costa Rica." *Revista de Biología Tropical* 36 (2A): 191–207.

Harvey, E. S., J. J. Butler, D. L. McLean, and J. Shand. 2012. "Contrasting Habitat Use of Diurnal and Nocturnal Fish Assemblages in Temperate Western Australia." *Journal of Experimental Marine Biology and Ecology* 426–427 (September): 78–86. https://doi.org/10.1016/j.jembe.2012.05.019.

Hiatt, Robert W., and Donald W. Strasburg. 1960. "Ecological Relationships of the Fish Fauna on Coral Reefs of the Marshall Islands." *Ecological Monographs* 30 (1): 65–127. https://doi.org/10.2307/1942181.

Hobson, Edmund S. 1965. "Diurnal – Nocturnal Activity of Some Inshore Fishes in the Gulf of California." *Copeia* 3: 291–302.

———. 1974. "Feeding Relationships of Teleostean Fishes on Coral Reefs in Kona, Hawaii" 72: 915–1031.

Horinouchi, Masahiro, Naoki Mizuno, Yuka Jo, Masashi Fujita, Yuzuru Suzuki, Futoshi Aranishi, and Mitsuhiko Sano. 2013. "Habitat Preference Rather than Predation Risk Determines the Distribution Patterns of Filefish Rudarius Ercodes in and around Seagrass Habitats." *Marine Ecology Progress Series* 488 (August): 255–66. https://doi.org/10.3354/meps10396.

Horinouchi, Masahiro, Mitsuhiko Sano, Toru Taniuchi, and Makoto Shimizu. 1998. "Food and Microhabitat Resource Use by Rudarius Ercodes and Ditrema Temmincki Coexisting in a Zostera Bed at Aburatsubo, Central Japan." *Fisheries Science* 64 (August): 563–68. https://doi.org/10.2331/fishsci.64.563.

Horinouchi, Masahiro, Prasert Tongnunui, Keisuke Furumitsu, Yohei Nakamura, Kouki Kanou, Atsuko Yamaguchi, Ken Okamoto, and Mitsuhiko Sano. 2012. "Food Habits of Small Fishes in Seagrass Habitats in Trang, Southern Thailand." *Fisheries Science* 78 (3): 577–87. https://doi.org/10.1007/s12562-012-0485-5.

Hutchins, J. Barry. 1997. *Review of the Monacanthid Fish Genus Paramonacanthus, with Descriptions of Three New Species*. Records of the Western Australian Museum, no. 54. Perth: Western Australian Museum.

Hutchins, J. Barry, and John E. Randall. 1982. "Cantherhines Longicaudaus, A New Filefish from Oceania, with a Review of the Species of the C. Fronticinctus Complex." *Pacific Science* 36 (2): 175–85.

Hyndes, G. A., A. J. Kendrick, L. D. MacArthur, and E. Stewart. 2003. "Differences in the Species- and Size-Composition of Fish Assemblages in Three Distinct Seagrass Habitats with

Differing Plant and Meadow Structure." *Marine Biology* 142 (6): 1195–1206. https://doi.org/10.1007/s00227-003-1010-2.
Jones, G. P. 1992. "Interactions between Herbivorous Fishes and Macro-Algae on a Temperate Rocky Reef." *Journal of Experimental Marine Biology and Ecology* 159 (2): 217–35. https://doi.org/10.1016/0022-0981(92)90038-C.
Kawase, Hiroshi. 2003. "Spawning Behavior and Biparental Egg Care of the Crosshatch Triggerfish, Xanthichthys Mento (Balistidae)." *Environmental Biology of Fishes* 66 (3): 211–19. https://doi.org/10.1023/A:1023978722744.
———. 2005. "Spawning Behavior of the Pygmy Leatherjacket Brachaluteres Jacksonianus (Monacanthidae) in Southeastern Australia." *Ichthyological Research* 52 (2): 194–97. https://doi.org/10.1007/s10228-005-0272-8.
Kawase, Hiroshi, and Akinobu Nakazono. 1994. "Reproductive Behavior of the Honeycomb Leatherjacket, Cantherhines Pardalis (Monacanthidae), at Kashiwajima, Japan" 41 (1): 80–83.
———. 1995. "Predominant Maternal Egg Care and Promiscuous Mating System in the Japanese Filefish, Rudarius Ercodes (Monacanthidae)." *Environmental Biology of Fishes* 43 (3): 241–54. https://doi.org/10.1007/BF00005856.
———. 1996. "Two Alternative Female Tactics in the Polygynous Mating System of the Threadsail Filefish,Stephanolepis Cirrhifer (Monacanthidae)." *Ichthyological Research* 43 (3): 315–23. https://doi.org/10.1007/BF02347603.
Khalaf, Maroof, and Uwe Zajonz. 2007. "Fourteen Additional Fish Species Recorded from below 150 m Depth in the Gulf of Aqaba, Including Liopropoma Lunulatum (Pisces: Serranidae), New Record for the Red Sea" 23: 421–33.
Kikuchi, Taiji. 1974. "Japanese Contributions on Consumer Ecology in Eelgrass (Zostera Marina L.) Beds, with Special Reference to Trophic Relationships and Resources in Inshore Fisheries." *Aquaculture* 4 (January): 145–60. https://doi.org/10.1016/0044-8486(74)90030-1.
Kim, Hye Rim, Jung Hwa Choi, and Won Gyu Park. 2013. "Vertical Distribution and Feeding Ecology of the Black Scraper, Thamnaconus Modestus, in the Southern Sea of Korea." *Turkish Journal of Fisheries and Aquatic Sciences* 13 (2): 249–59. https://doi.org/10.4194/1303-2712-v13_2_07.
Kuiter, Rudie H. 1992. *Tropical Reef-Fishes of the Western Pasific Indonesia and Adjacent Waters*. Indonesia: Penerbit Gramedia Pustaka Utama.
———. 2005. *Photo Guide to Fishes of the Maldives*. Apollo Bay, Vic.; Garsington: Atoll ; Windsor [distributor.
Kulbicki, Michel, Yves-Marie Bozec, Pierre Labrosse, Yves Letourneur, Gérard Mou-Tham, and Laurent Wantiez. 2005. "Diet Composition of Carnivorous Fishes from Coral Reef Lagoons of New Caledonia." *Aquatic Living Resources* 18 (3): 231–50. https://doi.org/10.1051/alr:2005029.
Kwak, Seok Nam, G. W. Baeck, and S Huh. 2003. "Feeding Habits of Stephanolepis Cirrifer in a Zostera Marina Bed." *Korean Journal of Ichthyology* 15 (4): 219–23.
Kwak, Seok Nam, Sung-Hoi Huh, and Chang Geun Choi. 2006. "Diel Changes in the Diet of Rudarius Ercodes: A Diurnal Omnivore and Nocturnal Carnivore." *Korean Journal of Ichthyology* 18 (3): 178–83.

Last, P. 1975. “Aspects of the Taxonomy and Ecology of Tasmanian Leatherjackets (Family Monacanthidae, Pisces).” Bachelor’s of Science (Honors), Hobart, Australia: University of Tasmania.

Layman, Craig A., and Brian R. Silliman. 2002. “Preliminary Survey and Diet Analysis of Juvenile Fishes of an Estuarine Creek on Andros Island, Bahamas.” *Bulletin of Marine Science* 70 (1): 199–210.

Lecchini, D., and R. Galzin. 2005. “Spatial Repartition and Ontogenetic Shifts in Habitat Use by Coral Reef Fishes (Moorea, French Polynesia).” *Marine Biology* 147 (1): 47–58. https://doi.org/10.1007/s00227-004-1543-z.

Lieske, Ewald, and Robert F. Myers. 2002. *Coral Reef Fishes. Indo-Pacific and Caribbean*. Rev. ed. Princeton Pocket Guides. Princeton, N.J: Princeton University Press.

Lindholm, R. 1984. “Observations on the Chinaman Leatherjacket Nelusetta Ayraudi (Quoy & Gaimard) in the Great Australian Bight.” *Marine and Freshwater Research* 35 (5): 597–99. https://doi.org/10.1071/mf9840597.

Livingston, Robert. 1982. “Trophic Organization of Fishes in a Coastal Seagrass System.” *Marine Ecology - Progressive Series* 7: 1–12. https://doi.org/10.3354/meps007001.

Longley, William Harding. 1941. *Systematic Catalogue of the Fishes of Tortugas, Florida, with Observations on Color, Habits, and Local Distribution*. Edited by Samuel F. Hildebrand. Vol. 34. Papers from Tortugas Laboratory. Washington, D.C.: Carnegie Institution of Washington. https://catalog.hathitrust.org/Record/001500824.

Lopez-Peralta, R. H., and C. A. T. Arcila. 2002. “Diet Composition of Fish Species from the Southern Continental Shelf of Colombia.” *Naga, Worldfish Center Quarterly* 25 (3–4): 23–29.

Mancera-Rodríguez, Néstor Javier, and José Juan Castro-Hernández. 2015. “Feeding Ecology of the Planehead Filefish Stephanolepis Hispidus (Pisces: Monacanthidae), in the Canary Islands Area.” *Revista de Biología Marina y Oceanografía* 50 (2): 221–34. https://doi.org/10.4067/S0718-19572015000300002.

Masuda, Reiji, Yoh Yamashita, and Michiya Matsuyama. 2008. “Jack Mackerel Trachurus Japonicus Juveniles Use Jellyfish for Predator Avoidance and as a Prey Collector.” *Fisheries Science* 74 (2): 276–84. https://doi.org/10.1111/j.1444-2906.2008.01522.x.

Matsuura, K. 2001. “Balistidae.” In *FAO Species Identification Guide for Fishery Purposes. The Living Marine Resources of the Western Central Pacific.*, 6. Bony fishes part 4. (Labridae to Latimeriidae), estuarine crocodiles.:3911–28. Rome, Italy: FAO.

Matsuura, K, and J. C. Tyler. 1997. “Tetraodontiform fishes, mostly from deep waters, of New Caldonia.” In *Résultats des campagnes MUSORSTOM*, edited by Bernard Séret, 17:173–208. Mémoires du Muséum National d’Histoire Naturelle Zoologie 174. Paris: Éd. du Muséum.

Matsuura, Keiichi, and Yoshihisa Shiobara. 1989. “A New Triggerfish, Rhinecanthus Abyssus , from the Ryukyu Islands.” *Japanese Journal of Ichthyology* 36 (3): 315–17. https://doi.org/10.1007/BF02905615.

McClanahan, T. R., and S. H. Shafir. 1990. “Causes and Consequences of Sea Urchin Abundance and Diversity in Kenyan Coral Reef Lagoons.” *Oecologia* 83 (3): 362–70. https://doi.org/10.1007/BF00317561.

Meyer, David L. 1985. “Evolutionary Implications of Predation on Recent Comatulid Crinoids from the Great Barrier Reef.” *Paleobiology* 11 (2): 154–64. https://doi.org/10.1017/S0094837300011477.

Myers, Robert F. 1991. *Micronesian Reef Fishes: A Practical Guide to the Identification of the Coral Reef Fishes of the Tropical Central and Western Pacific*. Barrigada, Territory of Guam, U.S.A.: Coral Graphics.

Nakamura, Yohei, Masahiro Horinouchi, Tadaomi Nakai, and Mitsuhiko Sano. 2003. “Food Habits of Fishes in a Seagrass Bed on a Fringing Coral Reef at Iriomote Island, Southern Japan.” *Ichthyological Research* 50 (1): 0015–0022. https://doi.org/10.1007/s102280300002.

Nakamura, Yohei, and Mitsuhiko Sano. 2004. “Overlaps in Habitat Use of Fishes between a Seagrass Bed and Adjacent Coral and Sand Areas at Amitori Bay, Iriomote Island, Japan: Importance of the Seagrass Bed as Juvenile Habitat.” *Fisheries Science* 70 (5): 788–803. https://doi.org/10.1111/j.1444-2906.2004.00872.x.

Patankar, Vardhan, Akhila Paranjape, Zoya Tyabji, Tanmay Wagh, and Aniruddha Marathe. 2018. “Occurrence and Distribution of Tetraodontiform Fishes of the Andaman and Nicobar Islands, India.” *Check List* 14 (3): 529–37. https://doi.org/10.15560/14.3.579.

Peristiwady, T., and P. Geistdoerfer. 1991. “Biological Aspects OfMonacanthus Tomentosus (Monacanthidae) in the Seagrass Beds of Kotania Bay, West Seram, Moluccas, Indonesia.” *Marine Biology* 109 (1): 135–39. https://doi.org/10.1007/BF01320240.

Randall, John E. 1955. “Fishes of the Gilbert Islands.” http://repository.si.edu//handle/10088/6126.

———. 1964. “A Revision of the Filefish Genera Amanses and Cantherhines.” *Copeia* 1964 (2): 331–61. https://doi.org/10.2307/1441027.

Randall, John E. 1967. *Food Habits of Reef Fishes of the West Indies*. [Coral Gables, Fla.] : Institute of Marine Sciences, University of Miami. https://trove.nla.gov.au/version/17839510.

Randall, John E. 1980. “Survey of Ciguatera at Enewetak and Bikini, Marshall Islands, with Notes on the Systematics and Food Habits of Ciguatoxic Fishes.” *Fish. Bull.; (United States)* 78 (2): 201–49.

———. 1985. *Guide to Hawaiian Reef Fishes*. Newtown Square, PA : Kaneohe, HI: Harrowood Books ; copublished and distributed in Hawaii by Treasures of Nature.

———. 1995. *Coastal Fishes of Oman*. Honolulu: University of Hawaii Press.

———. 2007. *Reef and Shore Fishes of the Hawaiian Islands*. Honolulu: Sea Grant College Program, University of Hawaiʻi.

Randall, John E., Gerald R. Allen, and Roger C. Steene. 1990. *Fishes of the Great Barrier Reef and Coral Sea*. Honolulu: University of Hawaii Press.

Randall, John E., John L. Earle, Richard L. Pyle, James D. Parrish, and Therese Hayes. 1993. “Annotated Checklist of the Fishes of Midway Atoll, Northwestern Hawaiian Islands.” *Pacific Science* 47 (4): 356–400.

Randall, John E., and W. D. Hartman. 1968. “Sponge-Feeding Fishes of the West Indies.” *Marine Biology* 1 (3): 216–25. https://doi.org/10.1007/BF00347115.

Randall, John E., Keiichi Matsuura, and Akira Zama. 1978. “A Revision of the Triggerfish Genus Xanthichthys, with Description of a New Species.” *Bulletin of Marine Science* 28 (October): 688–706.

Rodgers, Giverny G., Adrian J. Linnane, and Charlie Huveneers. 2013. “Contrasting Diet of Two Temperate Reef Fish Species (Notolabrus Tetricus and Meuschenia Hippocrepis) as Determined from Commercial Rock Lobster Bycatch Samples.” *Transactions of the Royal Society of South Australia* 137 (1): 80–89. https://doi.org/10.1080/3721426.2013.10887173.

Sancho, Gorka, C. Petersen, and Phillip Lobel. 2000. “Predator-Prey Relations at a Spawning Aggregation Site of Coral Reef Fishes.” *Marine Ecology Progress Series* 203 (September): 275–88. https://doi.org/10.3354/meps203275.

Sandin, Stuart A., and Ivor Williams. 2010. “Trophic Classifications of Reef Fishes from the Tropical U.S. Pacific (Version 1.0).” *Scripps Institution of Oceanography Technical Report.*, August. https://escholarship.org/uc/item/5394f7m3#author.

Sano, Mitsuhiko, Makoto Shimizu, and Yukio Nose. 1984. *Food Habits of Teleostean Reef Fishes in Okinawa Island, Southern Japan*. The University Museum, The University of Tokyo, Bulletin No. 25. Tokyo: University of Tokyo Press.

Schroeder, Robert. 1980. *Philippine Shore Fishes of the Western Sulu Sea*. Manila, Philippines: Bureau Of Fisheries And Aquatic Resources and NMPC Books.

Scott, J K. 1981. “The Seagrass Fish Fauna of Geographe Bay, Western Australia.” *Journal of the Royal Society of Western Australia* 63 (4): 97–102.

Smith, J. L. B., Margaret Mary Smith, and Phillip C. Heemstra. 1986. *Smiths’ Sea Fishes*. 1st ed. Braamfontein, Johannesburg: Macmillan South Africa.

Smith, Timothy, Gregory Jenkins, and Neil Hutchinson. 2012. “Seagrass Edge Effects on Fish Assemblages in Deep and Shallow Habitats.” *Estuarine, Coastal and Shelf Science* 115 (December): 291–99. https://doi.org/10.1016/j.ecss.2012.09.013.

Springer, Victor, and Kenneth Woodburn. 1960. “An Ecological Study of the Fishes of the Tampa Bay Area.” 1. Professional Papers Series. St. Petersburg, FL: Florida State Board of Conservation - Marine Laboratory. http://www.nativefishlab.net/library/textpdf/11938.pdf.

Tabb, Durbin C., and Raymond B. Manning. 1961. “A Checklist of the Flora and Fauna of Northern Florida Bay and Adjacent Brackish Waters of the Florida Mainland Collected During the Period July, 1957 through September, 1960.” *Bulletin of Marine Science of the Gulf and Caribbean* 11 (4): 552–649.

Taquet, Marc, Alain Diringer, John E Randall, Jean-Claude Quéro, Institut de recherche pour le développement (France), and Indo-Pacific Fish Conference (10th : 2017 : Tahiti). 2017. *Fishes of the Indian Ocean and the Red Sea*.

Taquet, Marc, Gorka Sancho, Laurent Dagorn, Jean-Claude Gaertner, David Itano, Riaz Aumeeruddy, Bertrand Wendling, and Christophe Peignon. 2007. “Characterizing Fish Communities Associated with Drifting Fish Aggregating Devices (FADs) in the Western Indian Ocean Using Underwater Visual Surveys.” *Aquatic Living Resources* 20 (4): 331–41. https://doi.org/10.1051/alr:2008007.

Travers, Michael J., I. C Potter, K. R Clarke, S. J. Newman, and J. Barry Hutchins. 2010. “The Inshore Fish Faunas over Soft Substrates and Reefs on the Tropical West Coast of

Australia Differ and Change with Latitude and Bioregion." *Journal of Biogeography* 37: 148–69.

Turingan, Ralph G. 1994. "Ecomorphological Relationships among Caribbean Tetraodontiform Fishes." *Journal of Zoology* 233 (3): 493–521. https://doi.org/10.1111/j.1469-7998.1994.tb05279.x.

Turingan, Ralph G., Peter C. Wainwright, and Dannie A. Hensley. 1995. "Interpopulation Variation in Prey Use and Feeding Biomechanics in Caribbean Triggerfishes." *Oecologia* 102 (3): 296–304. https://doi.org/10.1007/BF00329796.

Valesini, F. J, I. C Potter, and K. R Clarke. 2004. "To What Extent Are the Fish Compositions at Nearshore Sites along a Heterogeneous Coast Related to Habitat Type?" *Estuarine, Coastal and Shelf Science* 60 (4): 737–54. https://doi.org/10.1016/j.ecss.2004.03.012.

Vose, Frederic E., and Walter E. Nelson. 1994. "Gray Triggerfish (Balistes Capriscus Gmelin) Feeding from Artificial and Natural Substrate in Shallow Atlantic Waters of Florida" 55 (2–3): 1316–23.

Wenner, Charles A. 1983. "Species Associations and Day-Night Variability of Trawl-Caught Fishes from the Inshore Sponge-Coral Habitat, South Atlantic Bight" 81 (3): 537–52.

Witman, Jon D., Franz Smith, and Mark Novak. 2017. "Experimental Demonstration of a Trophic Cascade in the Galápagos Rocky Subtidal: Effects of Consumer Identity and Behavior." *PLOS ONE* 12 (4): e0175705. https://doi.org/10.1371/journal.pone.0175705.

Wressnig, Anna, and David J. Booth. 2008. "Patterns of Seagrass Biomass Removal by Two Temperate Australian Fishes (Monacanthidae)." *Marine and Freshwater Research* 59 (5): 408–17.

Wulff, J. 1994. "Sponge Feeding by Caribbean Angelfishes, Trunkfishes, and Filefishes." In *Sponges in Time and Space*, edited by van Soest, van Kempen, and Braekman. Rotterdam: Balkema.
